# SHMT1 and SHMT2 inhibition disrupts metabolic adaptation in glioblastoma

**DOI:** 10.64898/2026.11.17.767126

**Authors:** Jan-Hendrik Schröder, Leander F. Harwart, Sven König, Tanja Buhlmann, Hans Urban, Florian Strassheimer, Ann-Christin C. Hau, Jan-Bela Weinem, Dmitry Namgaladze, Mathias Haag, Sylvia K. Neef, Filip Hanak, Thorsten Mosler, Matthias Schwab, Lisa Hahnefeld, Irmgard Tegeder, Michael W. Ronellenfitsch, Joachim P. Steinbach, Anna-Luisa Luger

## Abstract

Glioblastoma cells adapt to hypoxia and nutrient restriction through metabolic reprogramming. Serine hydroxymethyltransferases 1 and 2 (SHMT1, SHMT2) channel serine-derived one-carbon units into folate metabolism, linking biosynthesis, respiration and redox defence. Its contribution to tumour metabolic homeostasis and adaptation to hypoxia and nutrient restriction remains unclear. We combined dual pharmacological SHMT inhibition using (+)-SHIN1 in glioblastoma cell lines, including patient-derived glioma stem-like models, with stable SHMT2 knockdown in LNT-229 cells. Functional analyses assessed clonogenic survival, invasion, migration and susceptibility to metabolic stress. Bioenergetics were measured by extracellular flux analysis and oxygen depletion, redox state by NADPH/NADP assay, and molecular responses by liquid chromatography–mass spectrometry-based metabolomics and ^13^C_6_-glucose tracing and tandem mass tag proteomics. Formate supplementation tested the contribution of limited one-carbon availability, three-dimensional co-cultures with human induced pluripotent stem cell-derived embryoid bodies assessed tumour expansion, and *in silico* analyses of public glioblastoma datasets examined *SHMT* expression, regional distribution and associations with hypoxia-related genes. *SHMT2* was enriched in hypoxic and perinecrotic tumour regions and correlated with hypoxia-inducible factor-1α target genes. SHMT inhibition impaired clonogenic growth, reduced invasion and migration, limited tumour expansion in co-culture, and sensitised cells to death under combined hypoxia and glucose restriction. Under normoxia, it increased respiration and estimates of glycolytic and oxidative ATP production. In G55T2 cells, SHMT inhibition lowered intracellular and extracellular lactate and altered tricarboxylic acid cycle metabolite levels and ¹³C isotopologue signals. SHMT inhibition increased oxidative stress, reduced the NADPH/NADP ratio in a cell line-and condition-dependent manner, and reduced extracellular formate accumulation. In G55T2 cells, SHMT inhibition led to a depletion of glutathione. Formate supplementation most prominently restored invasion and partially reversed other antitumour effects of SHMT inhibition, with the extent of rescue varying between cell lines. Under combined stress, SHMT inhibition attenuated hypoxia-inducible factor-1α accumulation, an effect counteracted by formate supplementation. These findings link SHMT-dependent one-carbon metabolism to redox homeostasis, invasive behaviour and survival under metabolic stress in glioblastoma. SHMT inhibition increases respiratory activity under normoxia while compromising tolerance to combined oxygen and glucose restriction, identifying metabolic adaptation as a potential therapeutic vulnerability in glioblastoma.

## Introduction

Glioblastoma is characterised by diffuse brain infiltration and pronounced spatial heterogeneity. Its microenvironment comprises cellular tumour regions, perivascular niches, hypoxic and peri-necrotic areas, and an infiltrative margin extending into surrounding brain tissue.^1^ Abnormal vascular organisation creates heterogeneous oxygen and nutrient availability, while tumour cells at the invasive edge encounter a different tissue environment from those within the tumour core. These regional conditions impose distinct demands on growth, survival and invasion, requiring glioblastoma cells to adapt to their microenvironment. Metabolic reprogramming is a hallmark of cancer that enables tumour cells to meet the demands of energy production and biosynthesis.^2^ The Warburg effect describes enhanced glucose conversion to lactate despite oxygen availability, a phenotype associated with the metabolic requirements of proliferating cells. However, aerobic glycolysis does not preclude mitochondrial oxidation.^3^ Glioblastoma cells exhibit metabolic plasticity, adjusting glycolysis, mitochondrial oxidation, and alternative substrate use according to oxygen and nutrient availability.^4,5^ Oxygen-dependent redistribution of glucose flux between the pentose phosphate pathway and glycolysis has also been demonstrated in glioma stem-like cells.^6^ Such adaptation requires coordination of energy metabolism, biosynthesis and redox defence, processes linked by serine-dependent one-carbon (1C) metabolism.^7,8^ Serine hydroxymethyltransferases (SHMTs) provide an entry point for serine-derived 1C units into folate metabolism. Cytosolic SHMT1 and mitochondrial SHMT2 catalyse the reversible transfer of a 1C unit from serine to tetrahydrofolate, generating glycine and 5,10-methylenetetrahydrofolate. They supply 1C units to compartmentalised folate-dependent pathways, with mitochondrially generated formate supporting cytosolic nucleotide biosynthesis.^8^ Reversal of cytosolic 1C flux can compensate for loss of the mitochondrial pathway in some cellular contexts.^8^ Folate-dependent reactions contribute to the production of reduced nicotinamide adenine dinucleotide phosphate (NADPH), while glycine provides a substrate for glutathione synthesis.^7,8^ SHMT-dependent metabolism may therefore regulate biosynthetic capacity and redox homeostasis during microenvironmental adaptation. SHMT2 also connects serine metabolism to mitochondrial respiration, although the direction of this relationship depends on the cellular context. In LN-229 glioblastoma cells, genetic *SHMT2* knockdown increases oxygen consumption, supporting a model in which SHMT2 restricts pyruvate kinase M2 (PKM2) activity and favours survival under ischaemic conditions.^5^ In other cellular models, SHMT2 supplies 1C units required for mitochondrial transfer RNA modifications^9^ and initiator tRNA formylation,^10^ thereby supporting mitochondrial translation and respiratory chain function. SHMT2 loss in these models reduces respiration. In glioblastoma cells, *HOTAIRM1* knockdown reduces both *SHMT2* expression and mitochondrial respiration, although this indirect intervention does not isolate the contribution of SHMT2 to the respiratory response.^11^ How dual pharmacological SHMT1 and SHMT2 inhibition affects bioenergetics and redox balance across glioblastoma models, and how these responses relate to survival under combined oxygen and nutrient limitation, remains incompletely understood. Beyond its biosynthetic and respiratory functions, serine-dependent 1C metabolism may contribute to invasive behaviour. Cells can export formate produced in excess of biosynthetic requirements, a process termed *formate overflow*.^12^ Formate can promote invasion in glioblastoma and other cancer models.^13^ This provides a link between mitochondrial serine catabolism and tumour dissemination in surrounding tissue. However, the contributions of SHMT-dependent metabolism to invasion and migration, and the extent to which these behaviours depend on formate availability, remain incompletely understood. The folate-competitive inhibitor (+)-SHIN1 targets both SHMT1 and SHMT2, enabling simultaneous pharmacological disruption of cytosolic and mitochondrial serine metabolism.^14^ Sensitivity of lymphoma cells to SHMT inhibition *in vitro*^15^ and antileukaemic activity of SHIN2 in preclinical T-cell acute lymphoblastic leukaemia models provide a rationale for investigating SHMT inhibition in glioblastoma.^16^ Here, we combined dual pharmacological SHMT inhibition with *SHMT2* knockdown to investigate glioblastoma cell survival, growth, invasion and migration. We assessed bioenergetic and redox responses and susceptibility to cell death under combined hypoxia and glucose restriction. Metabolite profiling, proteomics and formate supplementation were used to examine the metabolic basis of these responses, while three-dimensional co-cultures assessed their relevance to tumour expansion. We find that SHMT inhibition increases respiration in glioblastoma cells under normoxia and shifts glioblastoma cells towards a more oxidative state under normoxia while compromising their capacity to withstand combined hypoxia and glucose restriction, suggesting a conditional dependence on serine-derived one-carbon metabolism during microenvironmental stress.

## Materials and methods

### Analysis of public glioblastoma datasets

Gene expression analyses were performed using the GlioVis platform (https://gliovis.bio; accessed 12 Sep 2025).^17^ The Cancer Genome Atlas glioblastoma dataset (TCGA-GBM; accession phs000178; platform HG-U133A; *n* = 528 glioblastoma, *n* = 10 normal brain)^18^ was used to compare SHMT1 and SHMT2 expression between glioblastoma and normal brain. The Chinese Glioma Genome Atlas (CGGA; dataset mRNAseq_693; accessed 6 Sep 2025; *n* =225 for expression analysis)^19^ was used for correlations. Regional expression within glioblastoma (*n* = 111) was assessed using the Ivy Glioblastoma Atlas Project (Ivy GAP; accessed 28 Jan 2025).^20^ Only histologically diagnosed primary tumours were included.

### Cell lines and culture conditions

LN-229 (American Type Culture Collection), LNT-229 (TP53 wildtype; kind gift of N. de Tribolet, Lausanne, Switzerland),^21,22^ G55T2 (kind gift of M. Westphal and K. Lamszus, Hamburg, Germany),^23^ and the patient-derived glioma stem-like lines NCH644 (Cytion CLS, Eppelheim, Germany) and BG7 (kindly provided by H. Miletic, Bergen, Norway) were used. LN-229 identity was confirmed by short tandem repeat analysis (Multiplexion, Heidelberg, Germany); all lines were routinely tested mycoplasma-negative. LN-229, LNT-229 and G55T2 were cultured in high-glucose Dulbecco’s modified Eagle medium (DMEM) with 10% fetal calf serum (FCS) and penicillin/streptomycin. NCH644 and BG7 were cultured as suspension spheres in Neurobasal A medium supplemented with glutamine, B27, EGF and bFGF and dissociated with Accutase at approximately 300 µm diameter (Supplementary methods). Cells were maintained at 37°C in a humidified atmosphere with 5% CO2.

### Induction of hypoxia and glucose restriction

Hypoxia (0.1% O_2_) was induced in an anaerobic cultivation system and validated by chemical oxygen indicators, as described previously.^24,25^ For nutrient restriction, cells were cultured in serum-free DMEM formulated without glucose, glutamine or phenol red and supplemented with 2 mM glucose and penicillin/streptomycin.

### Drug treatments

Cells were allowed to adhere or recover for 24 h before treatment. Adherent cells were washed with 1x PBS and incubated in serum-free DMEM medium or DMEM supplemented with 2% FCS containing the indicated concentrations of (+)-SHIN1 (MedChemExpress, Germany) and/or sodium formate (Thermo Fisher Scientific, Germany). Suspension cultures received drug treatments at 2x the concentrations. (+)-SHIN1 was dissolved in dimethyl sulfoxide (DMSO), and DMSO at the highest final concentration served as vehicle control.

### Genetic manipulation of cell lines

BG7 cells were stably labelled with turboGFP using the GIPZ lentiviral vector (Open Biosystems, USA), with lentiviral particles produced as described previously.^26^ Stable SHMT2 knockdown in LNT-229 cells was achieved with a pLKO.1 vector encoding a short hairpin RNA (shRNA) targeting SHMT2 (Addgene, USA, TRCN0000238795); a non-targeting pLKO.1 construct (Addgene, USA, #1864) served as control (NTsh). Clones were selected with puromycin and knockdown was confirmed by immunoblotting. Seeding densities of the sublines were matched by crystal violet staining (Supplementary methods).

### Viability, cell death and clonogenic survival

Cellular ATP content was measured as a surrogate of viable cell abundance using a luminescent assay (CellTiter-Glo, Promega, USA) after 96 h of treatment. Clonogenic survival was assessed by colony formation from 1 × 10^3^ cells per well over 6-14 days, followed by crystal violet staining and manual counting.^24^ Cell death was quantified by propidium iodide (PI) staining with flow cytometry and, in parallel, by lactate dehydrogenase (LDH) release.^27^

### Flow cytometric assessment of oxidative stress and mitochondria

Reactive oxygen species-associated, oxidant-sensitive fluorescence was assessed using 2’,7’-dichlorodihydrofluorescein diacetate (H2DCFDA), and mitochondrial mass and membrane potential using MitoTracker Green and MitoTracker Red (Thermo Fisher Scientific, USA). Cells were counterstained with PI, and the mean fluorescence intensity of the PI-negative population was analysed in FlowJo (BD Biosciences, v10.10.0, USA).

### Invasion and migration assays

Invasion was assessed in Matrigel-coated Transwell inserts with 8 µm pore size (Sarstedt, Germany), with 10% FCS in the lower chamber as chemoattractant. After 24 h, invaded cells were stained with crystal violet and quantified in five randomly selected fields per insert at 20x magnification, blinded to treatment. Migration was measured as gap closure in two-well culture inserts (Ibidi, Germany) in LN-229 cells pre-treated with mitomycin C to inhibit proliferation; the remaining gap area at 24 and 48 h was expressed relative to the area at 0 h. Images were analysed in Fiji (ImageJ, version 2.16.0/1.54p) using standardised macros.

### Immunoblotting

Whole-cell lysates (10 µg protein per lane) were separated by sodium dodecyl sulphate-polyacrylamide gel electrophoresis on 12% gels, transferred to nitrocellulose membranes and probed with antibodies against SHMT1, SHMT2 (both Sigma-Aldrich, Germany) and beta-actin as loading control (Abcam, UK). Signals were detected by enhanced chemiluminescence and quantified by densitometry in Image Lab (version 6.1; Bio-Rad Laboratories, USA).

### Bioenergetic measurements

Real-time oxygen consumption was recorded over 24 h in 24-well plates with integrated optical oxygen sensors overlaid with paraffin oil (OxoDish, PreSens, Germany), as described previously.^27^ Oxygen consumption rate (OCR) and extracellular acidification rate (ECAR) were measured on an XF96 extracellular flux analyser (Agilent) after 24 h of treatment, using sequential injections of oligomycin (2.5 µM), carbonyl cyanide m-chlorophenyl hydrazone (1 µM) and rotenone (2.5 µM) with antimycin A (10 µg/ml). Data were normalised to protein content.

### Redox, formate and enzyme activity assays

The NADPH/NADP+ ratio,^25^ intracellular and extracellular formate, and pyruvate kinase activity were determined after 8 h of treatment using commercial luminescent or colorimetric assays (Promega, USA; Abcam, UK) according to the manufacturers’ protocols, with concentrations calculated from standard curves (Supplementary methods).

### Metabolomics

Following 8 h of treatment, the cell culture medium was collected for extracellular lactate quantification using targeted LC-MS/MS metabolomics, as previously described.^28^ Intracellular metabolites were extracted from 1 x 10^6^ cells per condition using ice-cold 80% methanol and analysed by LC-QTOF-MS for the semi-quantitative profiling of sugar phosphates, amino acids, tricarboxylic acid (TCA) cycle intermediates, and energy metabolites as described.^29^ Detailed extraction procedures and analytical conditions are provided in the Supplementary Methods.

### ¹³C-glucose tracing of polar metabolites

G55T2 cells were seeded 24 h before start of treatment. After taking the baseline samples, cells were switched to glucose-free, 2%-FCS DMEM supplemented with 25 mM (4.5 g/l) 13C_6_-Glucose and 10 µM (+)-SHIN1 or the respective volume of DMSO (control). Cells were harvested at 2 h, 4 h, 8 h and 16 h by trypsinisation. Cell pellets of 300,000 cells were stored at-80 °C until extraction. Incorporation of 13C-glucose into polar metabolites was examined using an UHPLC-HRMS metabolomic screening method^30^ using a Vanquish Horizon UHPLC system coupled to an Orbitrap Exploris 480 mass spectrometer, described in detail in the Supplementary Methods and Protocol (Excel file). Metabolites were identified using Compound Discoverer (v. 3.3) and the mzCloud library. Isotopic exchanges were subsequently curated in Skyline version 26.1.0.266.^31^

### Proteomics

Cell pellets were lysed, digested with LysC and trypsin, labelled with tandem mass tags and fractionated before analysis on an Orbitrap Fusion Lumos coupled to an easy nLC 1200 (Thermo Fisher Scientific), using a synchronous precursor selection multi-notch MS3 method to minimise ratio compression.^32^ Raw data were processed in FragPipe v22.0 with MSFragger v4.3^33^ and Philosopher v5.1.0,^34^ searched against the human reference proteome (UniProt; 20,418 entries) at a false discovery rate below 1% for peptide spectrum matches and proteins, quantified with IonQuant v1.11.11^35^ and further processed in FragPipe Analyst.^36^Full sample preparation, acquisition and database search parameters are provided in the Supplementary methods.

### Tumour spheroid-embryoid body confrontation assay

Embryoid bodies (EBs) were generated from human induced pluripotent stem cells (hiPSC; DF-19-9-11T.H, WiCell Research Institute, USA) following the STEMdiff Cerebral Organoid workflow and cultured until day 5, by which time they reached approximately 500 µm in diameter (Supplementary methods).^37^ GFP-labelled BG7 spheroids (2 × 10^3^ cells per well, 24 h) were co-cultured with EBs for 8 h before treatment and maintained for 3 days in mTeSR1-based medium with FGF2 and EGF (20 ng/ml each). Tumour and EB areas were segmented in Fiji/ImageJ at defined time points. Tumour expansion was calculated as

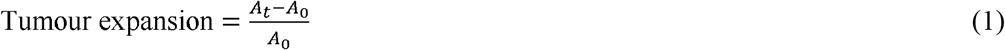

where A_t_ is the GFP-positive tumour area within the EB region of interest at time t and A0 the baseline area. The specific growth constant was calculated as

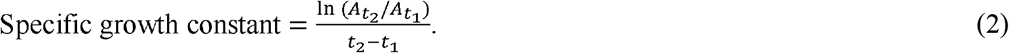

Bulk tumour area, with a minimum size threshold to exclude small satellite lesions, and EB area were quantified separately.

### Statistical analysis

Data are presented as mean ± standard deviation (SD) or standard error of the mean (SEM) unless otherwise specified in the figure legends. Analyses were performed in GraphPad Prism (version 10.5 or 11) and Microsoft Excel (version 16.113.2). Two groups were compared using a two-tailed Welch’s t-test. Three or more groups under a single factor were compared by one-way ANOVA, and experiments with two or more factors (e.g. time x treatment x oxygen) by multivariate ANOVA, each followed by Tukey’s or Dunnett’s multiple comparisons test as specified in the figure legends. Correlations were assessed using Spearman’s rank correlation coefficient. For oxygen consumption, slopes were calculated by linear regression analysis and tested as described above. *P* < 0.05 was considered statistically significant. Asterisks show adjusted P-values (\**P* < 0.05, \*\**P* < 0.01, \*\*\**P* < 0.001, \*\*\*\**P* < 0.0001).

## Results

### *SHMT* isoforms are differentially expressed and associated with adaptation to hypoxia in glioblastoma

We compared SHMT1 and SHMT2 mRNA expression in normal brain and glioblastoma using the TCGA-GBM dataset. *SHMT1* expression was significantly increased in glioblastoma compared to normal tissue, while *SHMT2* showed an even higher upregulation (Fig. 1A). SHMT1 and SHMT2 expression correlated significantly in the CGGA dataset (Fig. 1B), consistent with partially shared expression patterns. We assessed regional expression using the Ivy Glioblastoma Atlas Project (Ivy GAP) dataset. *SHMT1* is preferentially expressed in the cellular tumour regions at the infiltrative margin, including the leading edge, whereas *SHMT2* is enriched in hypoxia-and angiogenesis-associated regions, including microvascular proliferation, pseudopalisading cells around necrosis and perinecrotic zones (Fig. 1C). Consistent with its localization in hypoxic areas, *SHMT2* expression correlates significantly with multiple hypoxia-inducible factor 1α (HIF-1α; encoded by *HIF1A*) target genes, whereas *SHMT1* shows weaker correlations and no correlation with *VEGFA* (Fig. 1D). These analyses associate SHMT2 expression with hypoxia-related tumour regions in the analysed cohort.

**Figure 1.**
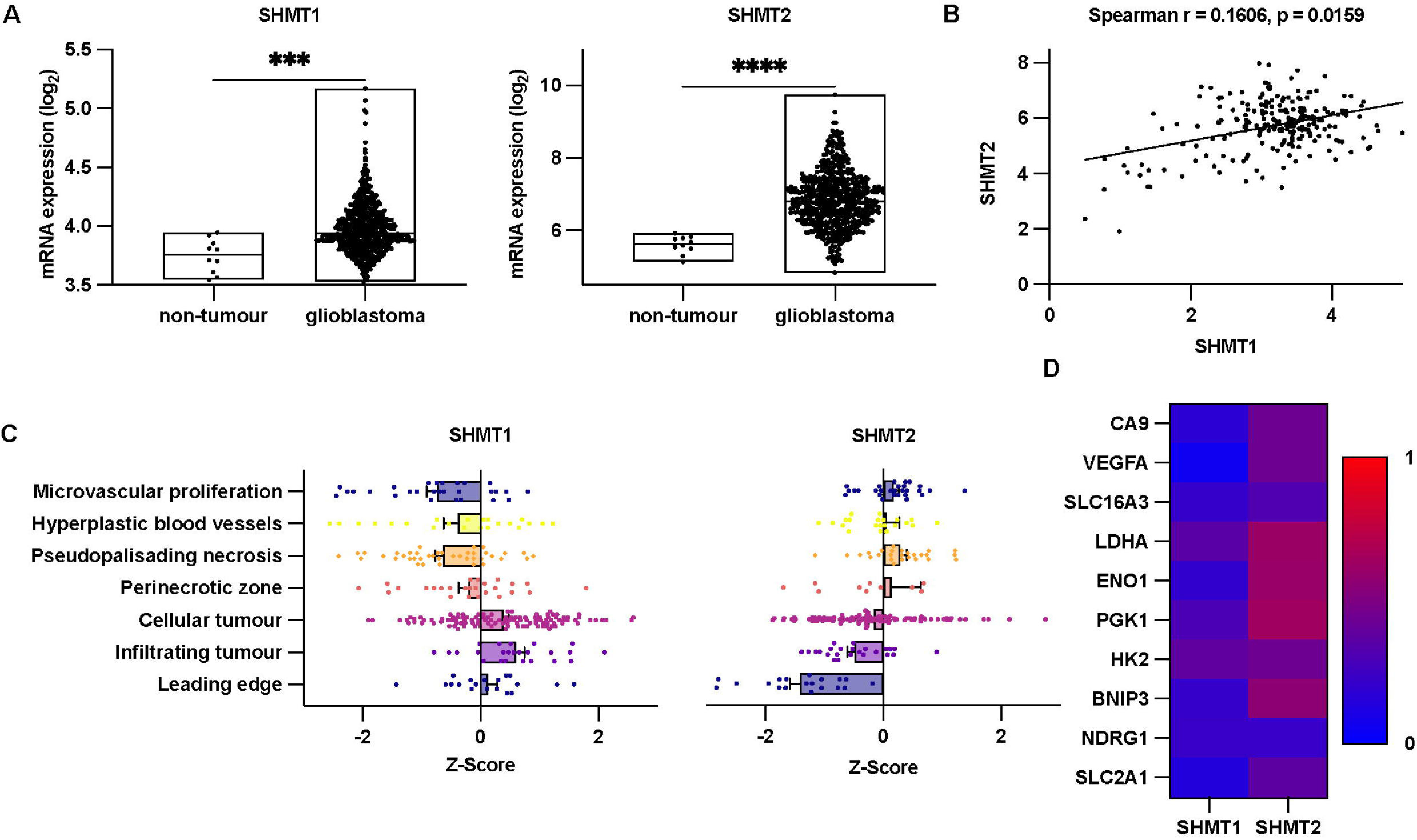
***SHMT* isoforms are differentially expressed and associated with adaptation to hypoxia in glioblastoma (A)** mRNA expression of *SHMT1* and *SHMT2* in glioblastoma (*n* = 528) versus normal brain tissue (*n* = 10) from the TCGA-GBM cohort analysed via GlioVis (median with minimum and maximum; two-tailed Welch’s t-test; \*\*\**P* < 0.001, \*\*\*\**P* < 0.0001). **(B)** Correlation of *SHMT1* and *SHMT2* transcript levels in glioblastoma using the CGGA database (*n* = 225; Spearman’s r). **(C)** Regional expression patterns of *SHMT1* and *SHMT2* in microvascular proliferation (*n* = 28), hyperplastic blood vessels (*n* = 22), pseudopalisading necrosis (*n* = 40), perinecrotic zone (*n* = 26), cellular tumour (*n* = 100), infiltrating tumour (*n* = 24), and leading edge (*n* = 19) based on Ivy GAP transcriptomic data (mean ± standard error of the mean). **(D)** Heatmap illustrating correlations of *SHMT1* and *SHMT2* expression with ten hypoxia-associated genes in the CGGA dataset (*n* = 225; Spearman’s r). Colour scale represents Spearman’s correlation coefficient (r), with red indicating strong positive correlation (r = 1) and blue indicating no correlation (r = 0).

### Dual SHMT1/2 inhibition reduces glioblastoma cell viability and clonogenicity and sensitises cells to hypoxia-induced death

We assessed cell viability after dual SHMT inhibition with (+)-SHIN1 in four glioblastoma cell lines. Viability decreased in all tested cell lines with mean half-maximal inhibitory concentration (IC) values ranging from 7.5 µM in LN-229 to 23.3 µM in NCH644 cells (Fig. 2A/B, S1A). Treatment of LN-229 cells with (+)-SHIN1 markedly reduced clonogenic growth in a concentration-dependent manner (Fig. 2C), supporting a role for SHMT activity in colony formation. To assess the relative importance of SHMT2, we established a stable *SHMT2* knockdown in LNT-229 cells using shRNA (Fig. S1B). SHMT1 levels were slightly increased, consistent with a potential compensatory upregulation (Fig. S1B). SHMT2 knockdown significantly reduced colony formation (Fig. 2D). We next examined whether SHMT inhibition sensitises glioblastoma cells to metabolic stress, resembling the hypoxic tumour microenvironment. Under combined hypoxia (0.1% O) and glucose restriction (2 mM), (+)-SHIN1 induced concentration-dependent cell death in LN-229 and NCH644 cells as assessed by propidium iodide (PI) staining and flow cytometry (Fig. 2E). Similarly, SHMT2 knockdown sensitised LNT-229 cells to combined hypoxia and glucose restriction supporting a contribution of SHMT2 to survival under metabolic stress (Fig. 2F). LDH-release assays further supported a concentration-dependent increase in cell death under combined hypoxia and glucose restriction following SHMT inhibition in LN-229 and G55T2 cells (Fig. S1C). Supplementation with 1 mM formate showed no changes in cell viability in NCH644 and BG7 cells (Fig. S1D), but partially rescued clonogenic growth in LN-229 and G55T2 cells (Fig. S1E). Formate also counteracted (+)-SHIN1-induced sensitisation to combined hypoxia and glucose restriction in LN-229 cells and, to a lesser extent, in G55T2 cells. No rescue was observed in BG7 cells, in which (+)-SHIN1 increased sensitivity to glucose restriction already under normoxia (Fig. S1F). Formate alone had no detectable effect on survival under combined hypoxia and glucose restriction in LN-229 and G55T2 cells (Fig. S1G). These findings support a role for SHMT activity in glioblastoma cell viability, clonogenic growth, and survival under metabolic stress. The variable rescue by formate suggests that impaired one-carbon supply contributes to these effects, with the extent of rescue depending on the model and experimental conditions.

**Figure 2.**
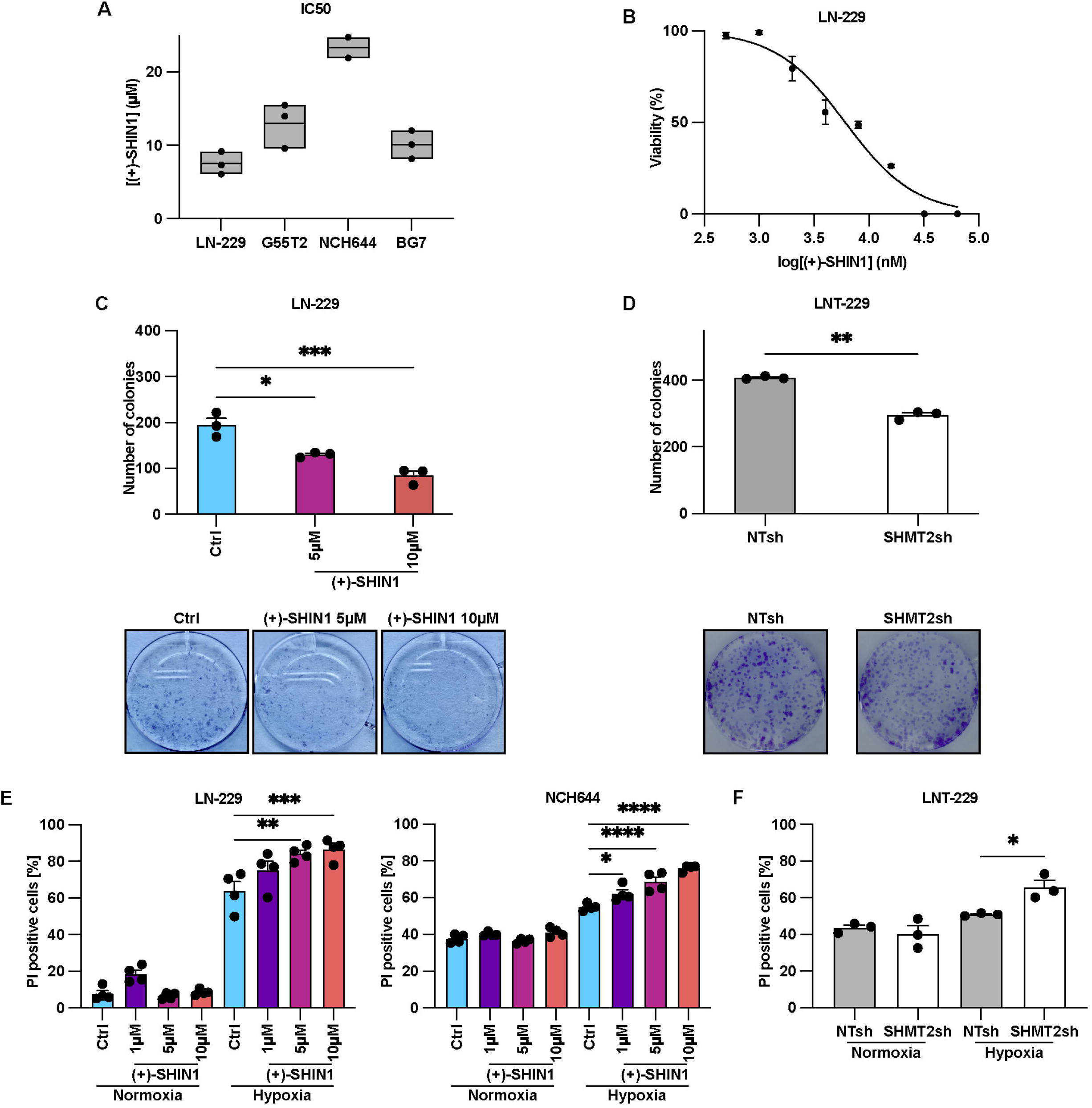
Dual SHMT1/2 inhibition reduces glioblastoma cell viability and clonogenicity and sensitises cells to hypoxia-induced death. **(A)** Half-maximal inhibitory concentration (IC_50_) values following (+)-SHIN1 treatment in glioma cell lines LN-229, G55T2, NCH644 and patient-derived glioma stem cells BG7 for 96 h in serum-free DMEM containing 25 mM glucose in normoxia (21% O_2_; *n* = 3; mean with maximum and minimum). **(B)** Cell viability of LN-229 cells following (+)-SHIN1 treatment in serum-free DMEM in normoxia (21% O_2_) for 96 h measured using the CellTiter-Glo assay (*n* = 3; mean ± standard error of the mean). **(C)** LN-229 were seeded (1,000 cells per well) in 6-well-plates, treated with 5 µM or 10 µM (+)-SHIN1 in DMEM containing 25 mM glucose and 10% fetal calf serum (FCS) in normoxia (21% O_2_) after 24 h and fixed by crystal-violet staining after 6 days (*n* = 3; mean; one-way ANOVA followed by Tukey’s multiple-comparisons test; \**P* < 0.05, \*\*\**P* < 0.001). Representative wells from one experiment are shown. **(D)** LNT-229 non-targeting shRNA (NTsh) and SHMT2-targeting shRNA (SHMT2sh) cells were seeded (1,000 cells per well) in 6-well-plates in DMEM containing 25 mM glucose and 10% FCS in normoxia (21% O_2_) and fixed by crystal-violet staining after 6 days (*n* = 3; mean; Welch’s t-test; \*\**P* < 0.01). Representative wells from one experiment are shown. **(E)-(F)** LN-229 and NCH644 treated with 1 µM, 5 µM or 10 µM (+)-SHIN1 and LNT-229 NTsh and SHMT2sh were incubated in serum-free DMEM containing 2 mM glucose in normoxia (21% O_2_) or hypoxia (0.1% O_2_) for 26-34 h. Cell death was analysed by propidium-iodide staining and quantified by flow cytometry (*n* = 4; mean ± standard error of the mean; one-way ANOVA followed by Tukey’s multiple-comparisons test; \**P* < 0.05, \*\**P* < 0.01, ***P < 0.001, \*\*\*\**P* < 0.0001). Vehicle controls received DMSO at the concentration present in the highest (+)-SHIN1 dose.

### SHMT inhibition suppresses glioblastoma cell invasion and migration

To evaluate the impact of SHMT inhibition on glioblastoma cell invasion, we performed Matrigel-coated Boyden chamber assays in four glioblastoma cell lines. Dual SHMT1 and SHMT2 inhibition with (+)-SHIN1 markedly reduced invasion in LN-229, G55T2, NCH644, and BG7 cells (Fig. 3A). Supplementation with 1 mM formate fully restored invasion in LN-229, G55T2, and NCH644 cells, whereas rescue was partial in BG7 cells (Fig. 3A). These data suggest that impaired folate-mediated one-carbon metabolism contributes to the reduced glioblastoma invasion upon SHMT inhibition. We assessed migration in the presence of mitomycin C to inhibit proliferation. In the gap-closure assay, (+)-SHIN1 significantly reduced gap closure in LN-229 cells (Fig. 3B). Formate supplementation only partially restored migration (Fig. 3B). We next examined invasion and gap closure in LNT-229 cells with stable SHMT2 knockdown. SHMT2 knockdown reduced invasion in the Boyden chamber assay, an effect at least partially reversed by formate supplementation (Fig. 3C). The effects of SHMT2 depletion in the gap closure assays were consistent with those in the Boyden chamber assay but overall weaker (Fig. 3D). Thus, SHMT depletion affects both invasion and gap closure, whereas formate supplementation preferentially rescues invasion under the conditions tested. To assess whether the two-dimensional effects of SHMT inhibition translate into a three-dimensional context, we employed a 3D confrontation assay combining embryoid bodies (EBs) with GFP-labelled BG7 tumour spheroids. Treatment with (+)-SHIN1 significantly reduced tumour expansion after two and three days (Fig. 3E). In this model, tumour expansion is contributed by growth and invasion. To assess bulk tumour growth separately, we analysed spheroid growth within the same co-culture system while excluding invasive outgrowth. (+)-SHIN1 treatment significantly reduced the specific growth constant (Fig. S2A), consistent with reduced bulk tumour growth in three-dimensional culture. In contrast, the specific growth constant of the embryoid bodies remained unchanged (Fig. S2B), suggesting a differential effect on tumour and embryoid body growth, without establishing tumour-selective toxicity. SHMT inhibition thus reduced both tumour expansion and bulk spheroid growth in three-dimensional co-culture.

**Figure 3.**
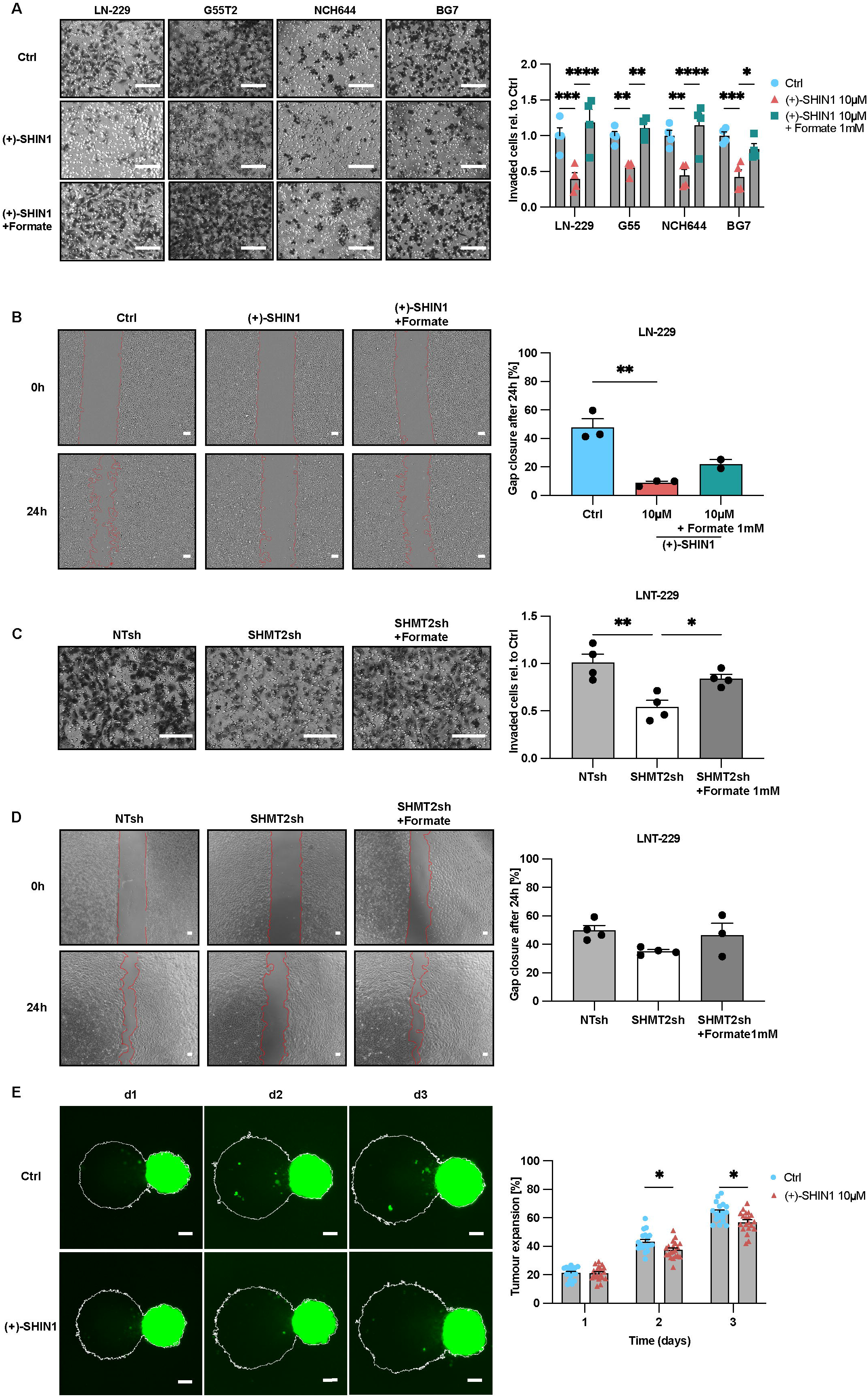
SHMT inhibition suppresses glioblastoma cell invasion and migration. **(A)** Invasion of glioma cell lines LN-229, G55T2, NCH644 and patient-derived glioma stem cells BG7 treated with 10 µM (+)-SHIN1 ± 1 mM formate in normoxia (21% O_2_) for 24 h in serum-free DMEM (n = 4; mean ± standard error of the mean; two-way ANOVA followed by Dunnett’s multiple-comparisons test; \**P* < 0.05, \*\**P* < 0.01, \*\*\**P* < 0.001, \*\*\*\**P* < 0.0001). Representative images from Matrigel-coated Boyden chambers. Scale bar = 150 µm. **(B)** Migratory potential of LN-229 cells in response to treatment in serum-free DMEM containing 25 mM glucose with 10 µM (+)-SHIN1 ± 1 mM formate in normoxia (21% O_2_) quantified as relative gap closure after 24 h (*n* = 3; mean ± standard error of the mean; one-way ANOVA followed by Tukey’s multiple-comparisons test; \*\**P* < 0.01). Representative images from the gap closure assay. Scale bar = 100 µm. **(C)** Invasion of LNT-229 non-targeting shRNA (NTsh) and SHMT2-targeting shRNA (SHMT2sh) cells incubated for 24 h in serum-free DMEM ± 1 mM formate in normoxia (21% O_2_; *n* = 4; mean ± standard error of the mean; one-way ANOVA followed by Tukey’s multiple-comparisons test; \**P* < 0.05, \*\**P* < 0.01). Representative images from Matrigel-coated Boyden chambers. Scale bar = 150 µm. **(D)** Migratory potential of LNT-229 NTsh and SHMT2sh cells incubated for 24 h in serum-free DMEM ± 1 mM formate in normoxia (21% O_2_; *n* = 4; mean ± standard error of the mean; one-way ANOVA followed by Tukey’s multiple-comparisons test). Representative images from the gap closure assay. Scale bar = 100 µm. **(E)** Quantitative representation of tumour expansion of GFP-labelled patient-derived glioma stem cells BG7 treated in mTeSR1-based medium with FGF2 and EGF (20 ng/ml each) with 10 µM (+)-SHIN1 in normoxia (21% O_2_) for 3 days co-cultured with embryoid bodies (EBs; *n* = 18-20 confrontation cultures; mean ± standard error of the mean; Welch’s t-tests followed by Holm–Šídák correction for multiple comparisons; \**P* < 0.05). Representative images from the co-culture invasion assay. Scale bar = 100 µm. Vehicle controls received DMSO at the concentration present in the highest (+)-SHIN1 dose.

### SHMT inhibition increases oxygen consumption and perturbs redox homeostasis

Given the role of folate-mediated one-carbon metabolism in cellular energy and redox balance, together with the differential rescue of the observed effects by formate, we next examined how SHMT inhibition affects bioenergetics and redox homeostasis in glioblastoma cells. In Seahorse analyses under normoxia, treatment with (+)-SHIN1 increased basal oxygen consumption rate (OCR) in both LN-229 and G55T2 cells (Fig. 4A). (+)-SHIN1 treatment increased oxygen depletion over 24 h in LN-229 and G55T2 cells in OxoDish measurements (Fig. S3A). Estimated glycolytic and oxidative ATP production also increased in both cell lines following treatment with 5 µM (+)-SHIN1 (Fig. 4B), consistent with increased bioenergetic activity under normoxia. Similarly, OxoDish measurements showed greater oxygen depletion over 24 h in SHMT2 knockdown LNT-229 cells, consistent with increased oxygen consumption (Fig. 4C). Additional treatment with (+)-SHIN1 did not further increase oxygen depletion in SHMT2-depleted cells (Fig. S3B). Formate supplementation attenuated the (+)-SHIN1-associated increase in OCR, whereas formate alone also increased OCR relative to untreated controls (Fig. S3C). (+)-SHIN1 treatment also increased extracellular acidification rate (ECAR). Formate supplementation attenuated this increase, whereas formate alone increased ECAR relative to untreated controls (Fig. S3D). We then assessed cellular redox homeostasis. H DCFDA fluorescence increased in a concentration-dependent manner in LN-229 cells under both normoxic and hypoxic conditions and in G55T2 selectively under hypoxia (Fig. 4D) and was also elevated in SHMT2-depleted LNT-229 cells under both conditions (Fig. 4E), consistent with increased intracellular oxidative stress. Formate alone did not significantly alter H DCFDA fluorescence (Fig. S3E) but significantly attenuated the (+)-SHIN1-induced increase under hypoxia in LN-229 cells (Fig. S3F). In G55T2 cells, the corresponding reduction was similar but weaker and did not reach statistical significance (Fig. S3F). We also assessed mitochondrial staining using MitoTracker Green as a proxy for mitochondrial mass and MitoTracker Red as a membrane potential-dependent mitochondrial stain. No significant treatment-associated changes in either MitoTracker signal were observed in LN-229 or G55T2 cells under normoxia or hypoxia (Fig. 4F). The increase in oxygen consumption therefore occurred without a detectable change in mitochondrial mass or membrane potential in these short-term experiments. To further characterise the redox alterations associated with SHMT inhibition, we next assessed the NADPH/NADP ratio. In LN-229 cells, the NADPH/NADP ratio declined significantly under both normoxia and hypoxia with glucose restriction, whereas NADPH/NADP^+^-ratio remained unchanged in G55T2 cells (Fig. 4G). Given the reported link between SHMT2 activity, PKM2 regulation, and oxygen consumption^5^ we also measured total pyruvate kinase activity. Pyruvate kinase activity was significantly increased by SHMT-inhibition under standard conditions in G55T2 cells, but not in LN-229 cells (Fig. 4 H). These findings show that SHMT inhibition increases basal respiration and estimated glycolytic and oxidative ATP production. These effects are accompanied by increased H DCFDA fluorescence and cell line-and condition-dependent reductions in the NADPH/NADP ratio.

**Figure 4.**
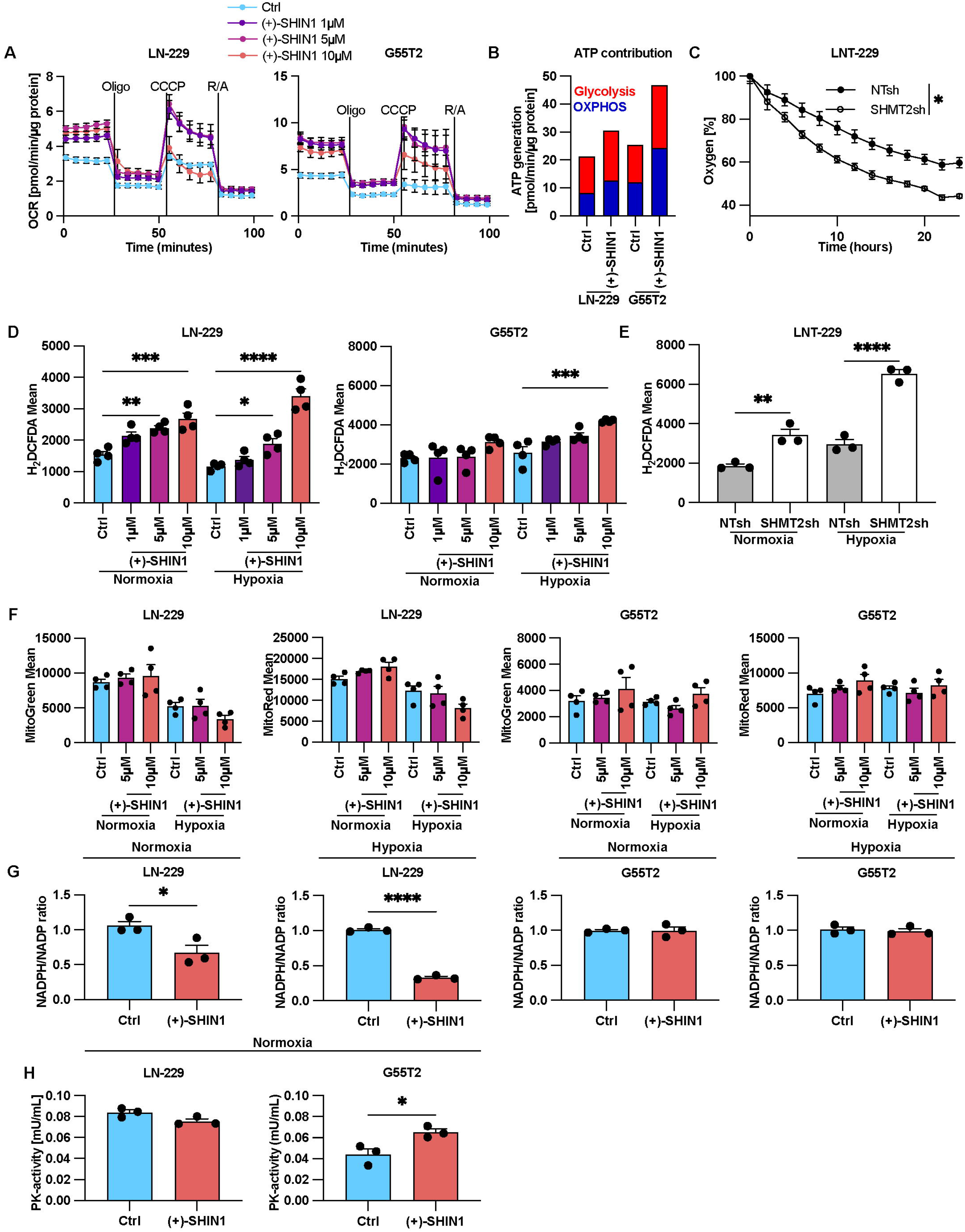
SHMT inhibition increases oxygen consumption and perturbs redox homeostasis. **(A)** LN-229 and G55T2 cells were incubated with 1 µM, 5 µM or 10 µM (+)-SHIN1 in serum-free DMEM containing 25 mM glucose for 24 h under normoxia (21% O_2_). Oxygen consumption rate (OCR) was measured in the Seahorse Bioscience Extracellular Flux Analyzer XF96 and XF^e^96 with port injections of oligomycin (Oligo), Carbonyl cyanide m-chlorophenyl hydrazone (CCCP), rotenone (R) and antimycin A (A) (*n* = 3; mean ± standard error of the mean). **(B)** Contribution of oxidative phosphorylation (OXPHOS) and glycolysis to ATP generation in LN-229 and G55T2 with or without (+)-SHIN1. **(C)** Oxygen consumption of LNT-229 non-targeting shRNA (NTsh) and SHMT2-targeting shRNA (SHMT2sh) cells was measured in DMEM containing 10% FCS in normoxia (21% O_2_) overlaid with paraffin oil with a fluorescence-based assay. Oxygen consumption is shown relative to the start of the experiment (*n* = 4; mean ± standard error of the mean; Welch’s t-test of linear regression slope; \**P* < 0.05). **(D)-(E)** LN-229 and G55T2 treated with 1 µM, 5 µM or 10 µM (+)-SHIN1 and LNT-229 NTsh and SHMT2sh cells were incubated in serum-free DMEM containing 2 mM glucose in normoxia (21% O_2_) or hypoxia (0.1% O_2_) for 18-24 h. Cells were analysed by 2′,7′-dichlorodihydrofluorescein diacetate (H2DCFDA) and propidium iodide (PI) staining. Only PI negative (= living) cells were quantified for H2DCFDA flow cytometry analysis (*n* = 4; mean ± standard error of the mean; one-way ANOVA followed by Tukey’s multiple-comparisons test; \**P* < 0.05, \*\**P* < 0.01, \*\*\*\**P* < 0.0001). **(F)** LN-229 and G55T2 cells were treated with 5 µM or 10 µM (+)-SHIN1 in serum-free DMEM containing 25 mM glucose in normoxia (21% O_2_) or 2 mM glucose in hypoxia (0.1% O_2_) for 8 h. Cells were analysed by MitoGreen, MitoRed and PI staining. Only PI negative (= living) cells were quantified for MitoTracker flow cytometry analysis (*n* = 4; mean ± standard error of the mean; one-way ANOVA followed by Tukey’s multiple-comparisons test). **(G)** LN-229 and G55T2 cells were treated with 10 µM (+)-SHIN1 in serum-free DMEM containing 25 mM glucose in normoxia (21% O_2_) or serum-free DMEM containing 2 mM glucose in hypoxia (0.1% O_2_) for 8 h. Analysis of NADPH/NADP^+^ ratios was performed by a luminescence-based assay (*n* = 3; mean ± standard error of the mean; Welch’s t-test; \**P* < 0.05, \*\*\*\**P* < 0.0001). **(H)** LN-229 and G55T2 cells were treated with 10 µM (+)-SHIN1 in serum-free DMEM containing 25 mM glucose in normoxia (21% O_2_) for 8 h. Pyruvate kinase activity was measured by a fluorescence-based assay kit (*n* = 3; mean ± standard error of the mean; Welch’s t-test; \**P* < 0.05, \*\**P* < 0.01, \*\*\*\**P* < 0.0001). Vehicle controls received DMSO at the concentration present in the highest (+)-SHIN1 dose.

### SHMT inhibition alters one-carbon metabolites, glutathione labelling and TCA cycle intermediates under normoxia

To assess the metabolic impact, we profiled intracellular metabolites and extracellular lactate in G55T2 cells after 8 h of treatment with 5 or 10 µM (+)-SHIN1 under normoxia (25 mM glucose) or combined hypoxia and glucose restriction (0.1% O, 2 mM glucose; Fig. 5A, B, C) and used ¹³C-glucose tracing to follow metabolite labelling under normoxia (Fig. 5D-H). Under normoxia, serine increased and glycine decreased, consistent with inhibition of SHMT-mediated serine-to-glycine conversion (Fig. 5A). (+)-SHIN1 effects on serine and glycine were less clear under hypoxia and glucose restriction, where both serine and glycine were highly increased as compared to normoxia. These stress conditions also caused a loss of intracellular formate, while extracellular formate accumulated (Fig. 5B, Fig S4A), the latter slightly attenuated in the presence of (+)-SHIN1, but drug effects were minor as compared to combined hypoxia and glucose restriction. Under normoxia, (+)-SHIN1 at both concentrations increased citrate and decreased succinate, malate and oxoglutaric acid. These TCA cycle metabolites showed no comparably pronounced, concentration-consistent changes under combined hypoxia and glucose restriction. In addition, intracellular and extracellular lactate decreased under normoxia (Fig. 5A, C), but remained fairly constant under combined hypoxia and glucose restriction (Fig. 5A, C). The changes in TCA cycle metabolite pools and lactate accompanied the increases in basal OCR and estimated oxidative ATP production described above (Fig. 4A, B). In the ¹³C-glucose time course we followed glutamate, which equilibrates rapidly with oxoglutaric acid, and fumarate, a key metabolic hub that links the TCA cycle with other pathways. For both, the abundance of both labelled and unlabelled isotopologues were lower after (+)-SHIN1 treatment at most time points (Fig. 5E, F). Above its functions as anaplerotic TCA metabolite, glutamate is an essential substrate for the first step of glutathione synthesis, while the second step essentially requires glycine. In line with that and the lower glycine levels, signals for ¹³C-labelled reduced and oxidised glutathione were lower after (+)-SHIN1 treatment (Fig. 5D). Branched chain amino acids including valine, leucine and isoleucine, which are essential amino acids, accumulated over the time course, consistent with reduced entry into branched-chain amino acid catabolism and/or increased release from protein turnover (Fig. 5G). Finally, we examined S-adenosylmethionine, the universal methyl donor and the second major output of the folate cycle alongside formate. After treatment with (+)-SHIN1, both unlabelled S-adenosylmethionine and its 13C-isotopologue were lower at 2 and 4 h and approached control values thereafter (Fig. 5H). Taken together, SHMT inhibition altered serine and glycine, lowered the incorporation of glucose-derived carbon into glutathione and reduced TCA cycle intermediates under normoxia. Both principal outputs of the folate cycle, formate and S-adenosylmethionine, were reduced.

**Figure 5.**
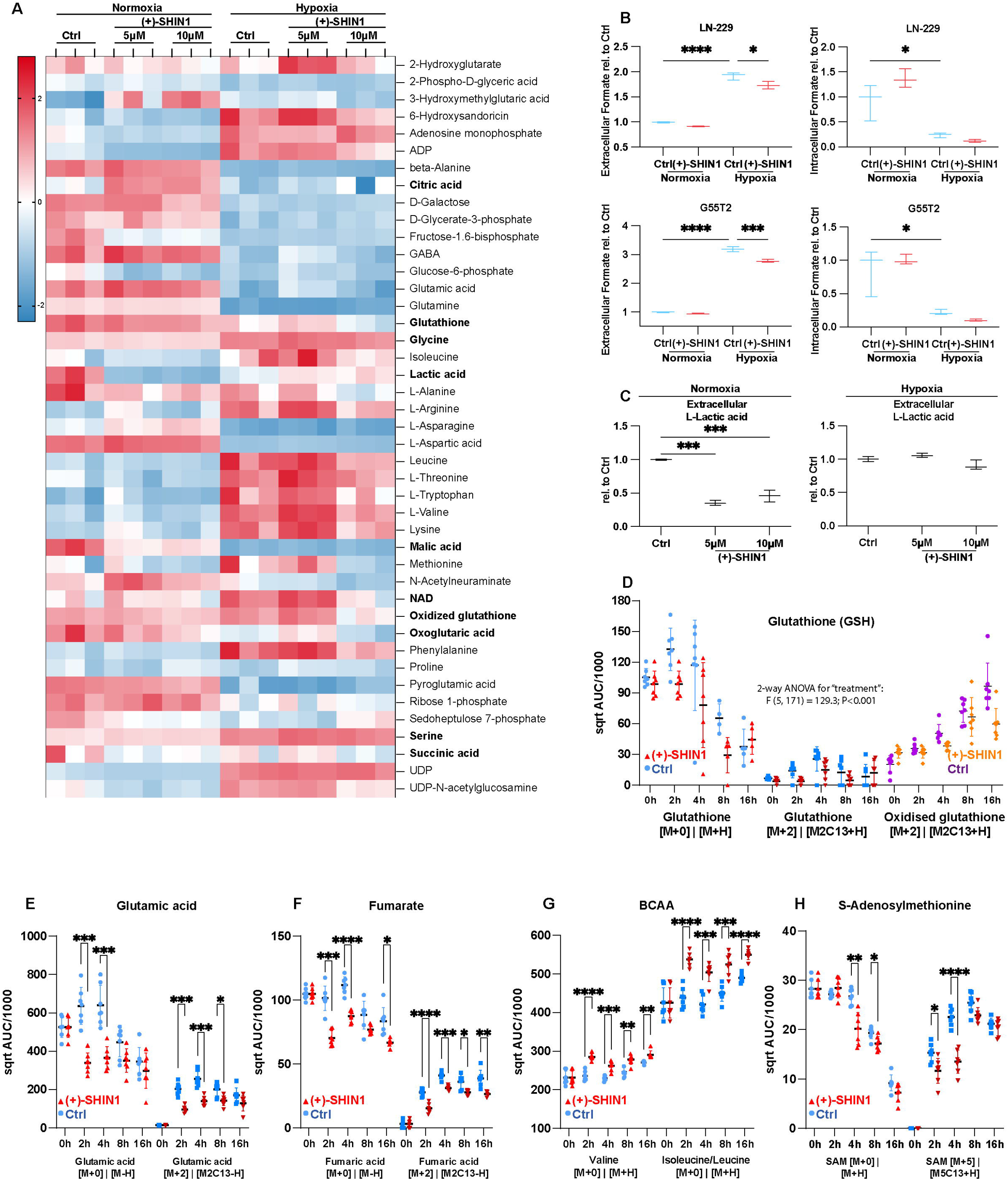
SHMT inhibition alters one-carbon metabolites, glutathione labelling and TCA cycle intermediates under normoxia. **(A)** Heatmap showing Z-score–normalised total peak areas of intracellular metabolites in G55T2 cells treated with 5 µM or 10 µM (+)-SHIN1 or vehicle control (Ctrl, DMSO concentration matching the highest (+)-SHIN1 concentration) for 8 h in serum-free DMEM containing 25 mM glucose in normoxia (21% O_2_) or 2 mM glucose in hypoxia (0.1% O_2_), as determined by untargeted LC-MS–based metabolite profiling. Each row shows a metabolite; each column shows a biological replicate. The colour scale shows the Z-score range, with red indicating higher and blue indicating lower relative abundance compared to the median across all conditions (*n* = 3 biological replicates). **(B)** Extra-and intracellular formate levels in LN-229 and G55T2 cells treated with 10 µM (+)-SHIN1 in serum-free DMEM containing 25 mM glucose in normoxia (21% O_2_) or 2 mM glucose in hypoxia (0.1% O_2_) for 8 h (*n* = 3; median with maximum and minimum; one-way ANOVA followed by Tukey’s multiple-comparisons test; \**P* < 0.05, \*\*\**P* < 0.001, \*\*\*\**P* < 0.0001). **(C)** Extracellular lactate levels with or without 5 µM or 10 µM (+)-SHIN1 in serum-free DMEM containing 25 mM glucose in normoxia (21% O_2_) or 2 mM glucose in hypoxia (0.1% O_2_; *n* = 3; median with minimum and maximum; one-way ANOVA followed by Tukey’s multiple-comparisons test; \*\**P* < 0.01, \*\*\**P* < 0.001). (**D**) Time course of glutathione (GSH, its heavy isotope), [M2C13+H], and oxidised GSH [M2C13+H] in G55T2 cells treated with 10 µM (+)-SHIN1 in serum-free DMEM containing 25 mM glucose in normoxia (21% O_2_; two-way ANOVA for “treatment”). **(E-H)** Time courses of glutamic acid, fumarate, branched chain amino acids (BCAA), and S-adenosylmethionine (SAM) in G55T2 cells treated with 10 µM (+)-SHIN1 in serum-free DMEM containing 25 mM glucose in normoxia (21% O_2_; two-way ANOVA “treatment” X “time followed by Tukey’s multiple-comparisons test; \**P* < 0.05, \*\**P* < 0.01, \*\*\**P* < 0.001, \*\*\*\**P* < 0.0001). Vehicle controls received DMSO at the concentration present in the highest (+)-SHIN1 dose.

### SHMT inhibition attenuates the hypoxic stress response and metabolic adaptation

To examine the consequences of SHMT inhibition on protein expression, we performed quantitative whole-cell proteomics in LN-229 cells under normoxia or combined hypoxia and glucose restriction (0.1% O, 2 mM glucose) (Fig. 6). In control cells, metabolic restriction increased the abundance of the hypoxia-associated proteins HIF1A and ANGPTL4 and the stress-response protein GDF15, as well as decreased the expression of the cholesterol biosynthesis-associated proteins squalene monooxygenase (SQLE) and 3-hydroxy, 3-methylglutaryl-CoA synthase (HMGCS) (Fig. S5A). Importantly, the defensive increase of HIF1A was reduced under (+)-SHIN1 treatment. Under normoxia, treatment with (+)-SHIN1 induced differential expression of proteins involved in mitochondrial metabolism, proliferation, DNA replication stress, and autophagy (Fig. 6A). Downregulated proteins included BCKDHB, the E1β subunit of the branched-chain ketoacid dehydrogenase complex that catalyses the committed step of branched-chain amino acid catabolism and the mitochondrial ribosomal proteins MRPL28 and MRPS11, consistent with changes in the mitochondrial translation machinery. Lower expression of proteins involved in proliferation and the upregulation of proteins that oppose DNA replication stress are consistent with constrained one-carbon supply and the observed reduction in proliferation.^38^ Under combined hypoxia and glucose restriction, (+)-SHIN1 reduced the expression of proteins associated with the hypoxia response, proliferation, mitochondrial biogenesis and the extracellular matrix, whereas proteins involved in glycolysis, mitochondrial metabolism, redox balance, and lipid metabolism were upregulated (Fig. 6B, 6C). In particular, the glioma invasion factor Tenascin-C (TNC) was downregulated, whereas citrate synthase (CS), hexokinase 1 (HK1), lactate dehydrogenases (LDHA/B), and the mitochondrial methionyl-tRNA formyltransferase (MTFMT) increased in abundance. The expression changes in redox-associated proteins agree with the observed condition-dependent alterations in cellular redox capacity, while decreased abundance of extracellular matrix and proliferation-associated proteins is in accordance with low invasion and clonogenic growth capacity under SHMT inhibition. Under combined metabolic stress, SHMT inhibition reduced the abundance of HIF-1α, the central coordinator of the transcriptional response to hypoxia (Fig. 6D). Supplementation with 1 mM formate significantly rescued this phenotype, supporting a contribution of SHMT-dependent one-carbon metabolism to the hypoxic response (Fig. 6D). These data associate SHMT inhibition with altered hypoxic stress response and adaptive metabolic reprogramming.

**Figure 6.**
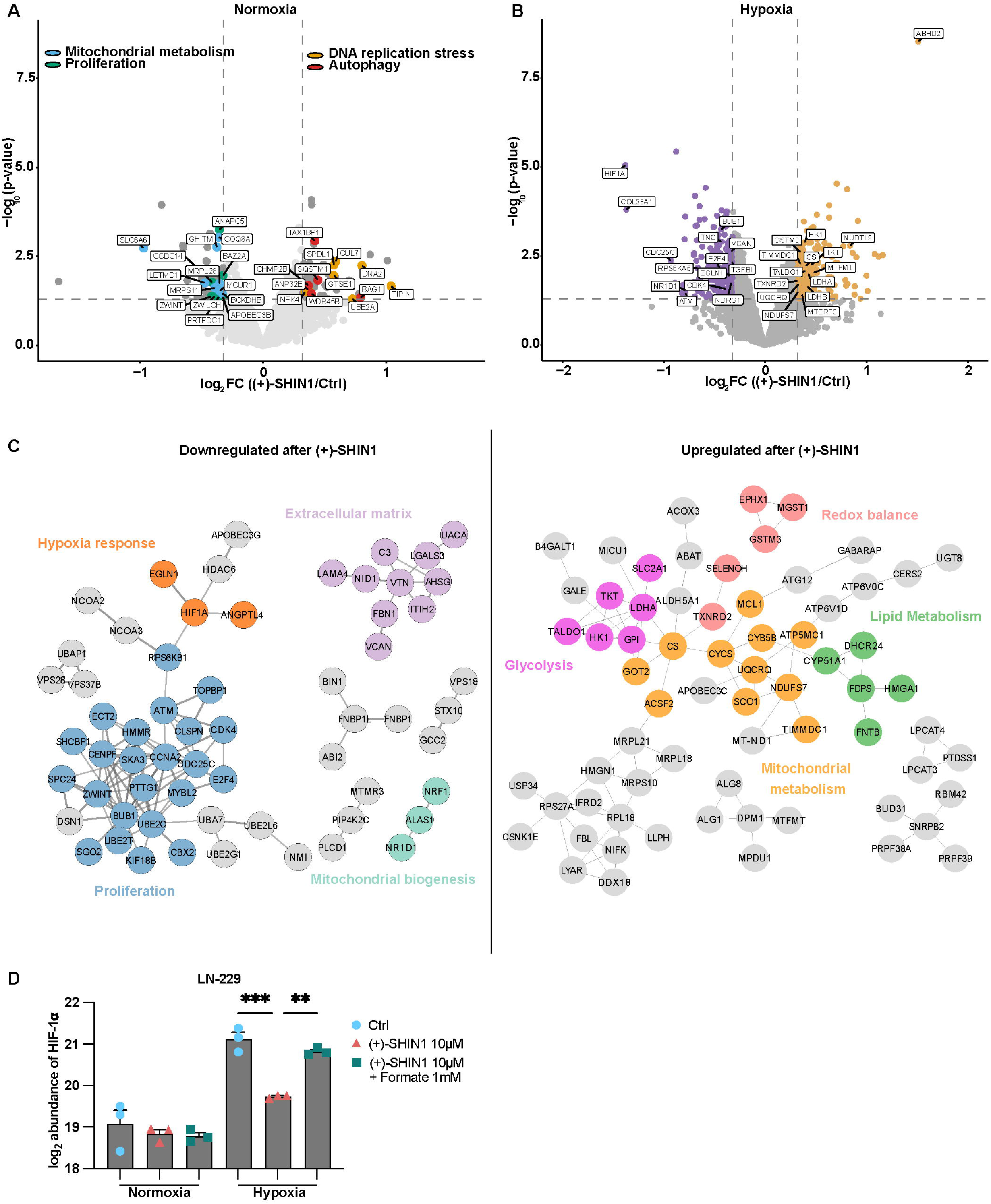
SHMT inhibition attenuates the hypoxic stress response and metabolic adaptation. **(A)** Volcano plot showing differential expression analysis of proteins in LN-229 cells treated with 10 µM (+)-SHIN1 versus vehicle in serum-free DMEM containing 25 mM glucose in normoxia (21% O_2_) for 8 h, analysed by tandem mass tag-based mass spectrometry. Proteins assigned to mitochondrial metabolism, proliferation, DNA replication stress and autophagy are highlighted. Dashed lines indicate the thresholds applied for significance (|log_2_ fold change| = 0.32, corresponding to a fold change of 1.25, and *P* = 0.05); the y-axis shows *P* values derived by limma^36,56^ (*n* = 3). **(B)** Corresponding comparison in serum-free DMEM containing 2 mM glucose in hypoxia (0.1% O_2_; *n* = 3). **(C)** Cytoscape^57,58^ protein–protein interaction networks of proteins downregulated (left) and upregulated (right) after (+)-SHIN1 under combined hypoxia and glucose restriction, with manual annotation of protein clusters colour-coded. **(D)** Bar plot displaying the log_2_ intensities of HIF-1α in LN-229 cells treated with 10 µM (+)-SHIN1 ± 1 mM formate in serum-free DMEM containing 25 mM glucose in normoxia (21% O_2_) or 2 mM glucose in hypoxia (0.1% O_2_; *n* = 3; mean ± standard error of the mean; one-way ANOVA followed by Tukey’s multiple-comparisons test; \*\**P* < 0.01, \*\*\**P* < 0.001). Vehicle controls received DMSO at the concentration present in the highest (+)-SHIN1 dose.

## Discussion

Inhibition of serine-dependent one-carbon metabolism with (+)-SHIN1 reorganises glioblastoma metabolism. By restricting the supply of one-carbon units from serine, (+)-SHIN1 may limit de novo purine and thymidylate synthesis, in analogy to classical antifolates such as methotrexate, which exploit nucleotide depletion to induce replication stress and cell cycle arrest.^38^ Consistent with this interpretation, proteins associated with DNA replication stress were among those altered by (+)-SHIN1 under normoxia, interfering thereby with several malignant phenotypes. Dual SHMT inhibition reduced metabolic viability in a panel of glioblastoma lines. Sub-IC concentrations of (+)-SHIN1 markedly impaired clonogenic survival, suggesting that SHMT inhibition at these concentrations undermines the long-term proliferative potential of individual tumour cells. Invasion and migration represent additional hallmarks that are tightly coupled to poor outcome in glioblastoma.^1,2,39^ In our Transwell assays, (+)-SHIN1 significantly reduced invasion across all tested glioblastoma cell models, and similarly, SHMT2 knockdown attenuated invasion. SHMT inhibition also increased sensitivity to cell death under hypoxia and glucose restriction. In a 3D co-culture system combining patient-derived BG7 spheroids with human iPSC-derived embryoid bodies, (+)-SHIN1 reduced tumour expansion and decreased the specific growth constant of the tumour spheroids without affecting embryoid body growth, confirming that SHMT inhibition disrupts glioblastoma expansion in a 3D organotypic context without establishing tumour-selective toxicity. The analyses of TCGA, CGGA and Ivy GAP datasets show an association of SHMT2 expression with hypoxia-related glioblastoma regions. Both SHMT isoforms are upregulated in glioblastoma relative to normal brain, but SHMT2 shows the strongest overexpression, correlates with hypoxia signatures and HIF-1α target genes, and is enriched in peri-necrotic, pseudopalisading and microvascular proliferation zones, in line with previous findings.^5^ These regions are characterised by extreme hypoxia, glucose restriction and therapy-resistant cell populations.^5,25,40,41^ By contrast, SHMT1 is preferentially expressed in infiltrative and leading-edge regions. The distinct spatial pattern and positive correlation between SHMT1 and SHMT2 expression point to complementary roles and coordinated regulation. SHMT inhibition directly affects the serine–glycine axis. By blocking the interconversion of serine and glycine, (+)-SHIN1 reduces metabolic flexibility in tumours known to be serine-and glycine-dependent.^14,42^ In G55T2 cells, serine accumulated and glycine levels decreased under normoxia, consistent with inhibition of the primary SHMT-mediated catalysis, the serine-to-glycine conversion. Importantly, this reaction also fuels the mitochondrial folate pathway via MTHFD2, which acts as a primary producer of mitochondrial NADPH ^7^, and provides glycine for glutathione production. Reduced availability of SHMT-derived glycine therefore contributes to the lowering of glutathione pools. A central mechanistic insight from our study is that SHMT inhibition reduces combined hypoxia and glucose restriction-evoked extracellular formate accumulation.^10,12,13,43,44^ In the “formate overflow” model, serine catabolism to formate allows tumours to maintain DNA synthesis and to gain one net ATP per exported formate molecule, thereby supporting growth and invasion.^12,45^ The observed rise of extracellular formate while depleting the intracellular pool under combined hypoxia and glucose restriction is consistent with altered formate handling under oxygen and glucose restriction, although pool measurements cannot differentiate between excessive formate production or efflux. The lowering of extracellular formate was most pronounced under combined hypoxia and glucose restriction, but (+)-SHIN1 treatment did not restore normoxic formate levels. Nonetheless, supplementation of exogenous formate restored invasiveness in (+)-SHIN1-treated cell lines and in SHMT2 knockdown cells and attenuated the loss of HIF-1α under metabolic stress while partially rescuing clonogenic growth and survival in a model-dependent manner. Hence, we conclude that even a moderate reduction of one-carbon availability in the tumour environment in combination with glycine and NADPH deficiency under (+)-SHIN1 treatment account for its anti-tumour efficacy. (+)-SHIN1 treatment also transiently reduced S-adenosylmethionine, a methyl donor for DNA and histone methylation. Whether this decrease had epigenetic consequences was not examined in the present study. Under normoxia, (+)-SHIN1 increased basal OCR and estimated oxidative ATP production, alongside an increase in estimated glycolytic ATP production. In G55T2 cells, intracellular and extracellular lactate decreased, and TCA cycle metabolite pools and ¹³C isotopologue signals changed. The pool sizes and isotopologue signals do not establish the direction of change in TCA cycle flux. The accumulation of BCAA reflects a shift between consumption and release. BCAA catabolism involves transamination with α-ketoglutarate and subsequent decarboxylation by the branched-chain ketoacid dehydrogenase complex (BCKDHB), which supplies succinyl-CoA and acetyl-CoA to the cycle. Since α-ketoglutarate was reduced and BCKDHB downregulated we infer that both steps of BCAA catabolism are constrained under SHMT inhibition in normoxia. An additional release of amino acids from protein turnover may contribute to the observed raise of BCAA, in line with the upregulation of autophagy-associated proteins under the same condition. The increase in respiration is consistent with previous observations in SHMT2-deficient tumour cells, in which increased oxidative metabolism has been linked to alterations in glycolytic control, including regulation of PKM2 by various metabolites including serine.^5,46–48^ Nevertheless, the idea of altered partitioning of glucose carbon between serine synthesis and mitochondrial metabolism remains a hypothesis.^49^ In our studies, total pyruvate kinase activity increased in G55T2 but not LN-229 cells. Our proteomic data suggest that (+)-SHIN1 impairs adaptation to combined hypoxia and glucose restriction. Under combined hypoxia and glucose restriction, SHMT inhibition attenuated the hypoxia evoked response, including the rise of HIF-1α itself, together with extracellular matrix constituents and cell cycle regulators, while inducing glycolytic, mitochondrial and redox proteins, such as GSTM3. The enrichment of seemingly opposing pathways may reflect compensatory mechanisms because for example, redox enzymes increased whereas enzymes involved in maintenance of the redox capacity decreased, and the mitochondrial methionyl-tRNA formyltransferase MTFMT, which formylates the mitochondrial initiator tRNA, also increased.^9,10^ Downregulated pathways were in line with the growth patterns, from reduced invasion to reduced clonogenic growth and viability. The comparison of normoxic and hypoxic metabolic profiles suggest that SHMT inhibition and combined hypoxia and glucose restriction share some metabolic consequences. Under normoxia, SHMT inhibition increased oxygen consumption and the estimated oxidative ATP production without detectable changes in mitochondrial mass or membrane potential (Fig. 4A, B, F). At the same time, a number of mitochondrial ribosomal proteins were downregulated under basal conditions. Under combined hypoxia and glucose restriction, reductions in redox capacity were cell line dependent and were accompanied by increased oxidant-sensitive fluorescence (Fig. 4D, G). We therefore propose that SHMT inhibition sensitises glioblastoma cells to the hypoxic, nutrient-poor microenvironment by altering metabolic demand and compromising redox homeostasis. In line with this reasoning, it has been reported that glioma cells in paediatric diffuse midline glioma models perform a metabolic shift, where ERK5 knockdown reduced extracellular lactate levels while oxygen consumption and expression of respiratory chain proteins were increased. In the same study, combined pharmacological inhibition of ERK5 and PFKFB3 (6-phosphofructo-2-kinase/fructose-2,6-bisphosphatase 3) increased ROS-associated fluorescence and cell death.^50^ The observation that HIF-1α levels dropped while oxidant-sensitive fluorescence increased is no contradiction.^51^ HIF-1α is short-lived and depends on continuous synthesis and ubiquitin-mediated degradation. HIF-1α stability is fuelled by one carbon metabolism, supported by our formate-rescue experiments, because 1C homeostasis provides the environment required for its stability. Inversely, HIF-1α drives 1C metabolism by maintaining transcription of crucial enzymes. Because hypoxia itself drives a mesenchymal, pro-invasive programme in glioblastoma,^40^ the loss of matrix proteins under these conditions rises the possibility that the observed anti-invasive effect under normoxia could also extend to hypoxic tumour regions. We did not test this directly, since (+)-SHIN1 induces cell death under combined hypoxia and glucose restriction which would confound an invasion readout. Dual SHMT inhibition may impair adaptation in hypoxic, therapy-resistant subpopulations that depend on serine-derived one-carbon metabolism for redox homeostasis and survival. These findings suggest combination strategies for further testing. The proposed anti-Warburg-like response and impaired redox buffering provide a rationale for testing combinations with radiotherapy, although increased oxygen consumption could equally aggravate hypoxia and radio-sensitisation.^52^ Restricting nucleotide pools and impairing redox buffering could also enhance the efficacy of temozolomide, whose cytotoxicity depends on DNA damage and limited repair capacity.^53^ In haematologic malignancies, antifolate strategies have already shown promise, providing a rationale to investigate whether related combinations are effective in glioblastoma.^15,16^ Such strategies are supported by our data because SHMT inhibition increased levels of proteins assigned to autophagy under normoxic conditions. In addition, dietary serine and glycine restriction has been shown to synergise with inhibition of serine synthesis in preclinical models,^42^ raising the possibility that metabolic interventions could potentiate pharmacological SHMT blockade in selected patients. We are now testing this hypothesis in clinical settings in the ongoing B-FREE window-of-opportunity trial (DRKS00038325),^54^ which assesses the metabolic and immunological effects of a normocaloric serine-and glycine-free diet in patients with glioma before surgery. Finally, SHMT2 enrichment in hypoxic regions rises the question of how SHMT inhibition would affect the immune microenvironment, given that serine metabolism is required for lymphocyte proliferation and effector function.^55^ Whether SHMT inhibition favours or impairs immune effector function requires direct testing *in vivo,* pointing to one of the limitations of our study. Further, the metabolic conclusions rest in part on steady-state metabolite levels and respirometry. Isotope tracing was performed in one cell line at one (+)-SHIN1 concentration. Therefore, metabolomics mainly describes altered pools and consumption rates while pathway flux was obtained for selected metabolites. All experiments were performed *in vitro* or in three-dimensional co-culture, and the therapeutic potential of SHMT inhibition in glioblastoma remains to be established *in vivo* using inhibitors with adequate brain penetration, alongside assessment of their effects on non-malignant brain cells and the immune response.

In line with previous studies, our data link formate availability to invasive cell features and survival under microenvironmental stress, and we provide evidence that SHMT inhibition may be an effective strategy to combat formate-dependent tumour resistance.

## Data availability

The datasets used and/or analysed in this study are available from the corresponding author upon reasonable request.

## Acknowledgements

We thank all members of the Centre for Functional Proteomics (Goethe University, Frankfurt). We thank the Deutsche Forschungsgemeinschaft (German Research Foundation, DFG) for funding the LC-MS system (easy nLC1200, Orbitrap Fusion LUMOS) used in this study (FuGG Project-ID: 403765277). Markus König is gratefully acknowledged for his technical assistance with the metabolomics analysis. We thank Carlo Angioni (Institute of Clinical Pharmacology, Goethe-University, Frankfurt, Germany) for his technical support and assistance in UHPLC-HRMS measurements. The authors used Claude (Anthropic) and ChatGPT (OpenAI) for language editing and for checking the internal consistency of the manuscript. These tools were not used to generate or analyse data, or to draft scientific content. The authors reviewed and verified all output and take full responsibility for the content of this manuscript. No artificial intelligence tool fulfils the criteria for authorship.

