## Supplementary material for "SHMT1 and SHMT2 inhibition disrupts metabolic adaptation in glioblastoma"

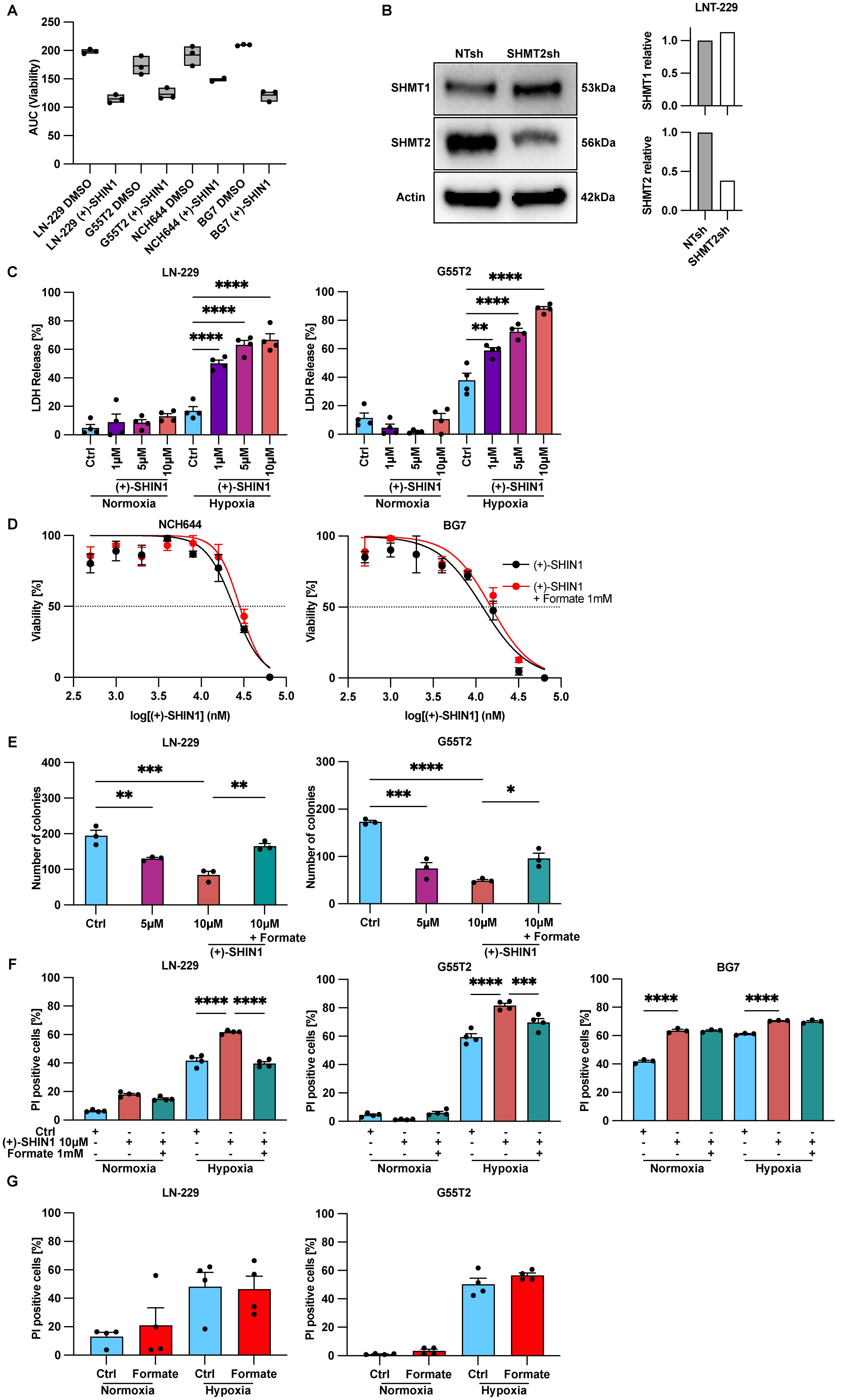

A
AUC (Viability)
LN-229 DMSO
LN-229 (+)-SHIN1
G55T2 DMSO
G55T2 (+)-SHIN1
NCH644 DMSO
NCH644 (+)-SHIN1
BG7 DMSO
BG7 (+)-SHIN1
B
NTsh
SHMT2sh
SHMT1
SHMT2
Actin
53kDa
56kDa
42kDa
LNT-229
SHMT1 relative
SHMT2 relative
NTsh
SHMT2sh
C
LN-229
G55T2
LDH Release [%]
Ctrl
1µM
5µM
10µM
(+)-SHIN1
Normoxia
Hypoxia
D
NCH644
BG7
Viability [%]
log[(+)-SHIN1] (nM)
(+)-SHIN1
(+)-SHIN1
+ Formate 1mM
E
LN-229
G55T2
Number of colonies
Ctrl
5µM
10µM
10µM
+ Formate
(+)-SHIN1
F
LN-229
G55T2
BG7
PI positive cells [%]
Ctrl
(+)-SHIN1 10µM
Formate 1mM
Normoxia
Hypoxia
G
LN-229
G55T2
PI positive cells [%]
Ctrl
Formate
Normoxia
Hypoxia

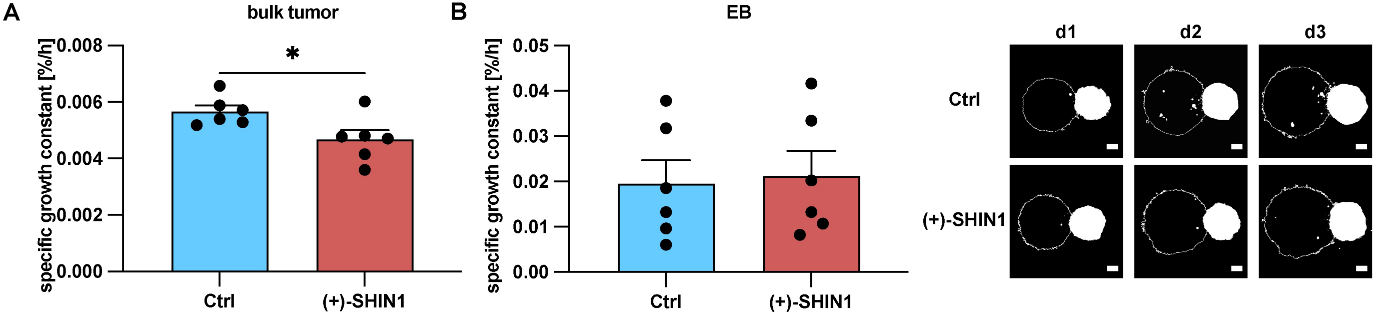

A
bulk tumor
specific growth constant [%/h]
0.008
0.006
0.004
0.002
0.000
*
Ctrl
(+)-SHIN1
B
EB
specific growth constant [%/h]
0.05
0.04
0.03
0.02
0.01
0.00
Ctrl
(+)-SHIN1
d1
d2
d3
Ctrl
(+)-SHIN1

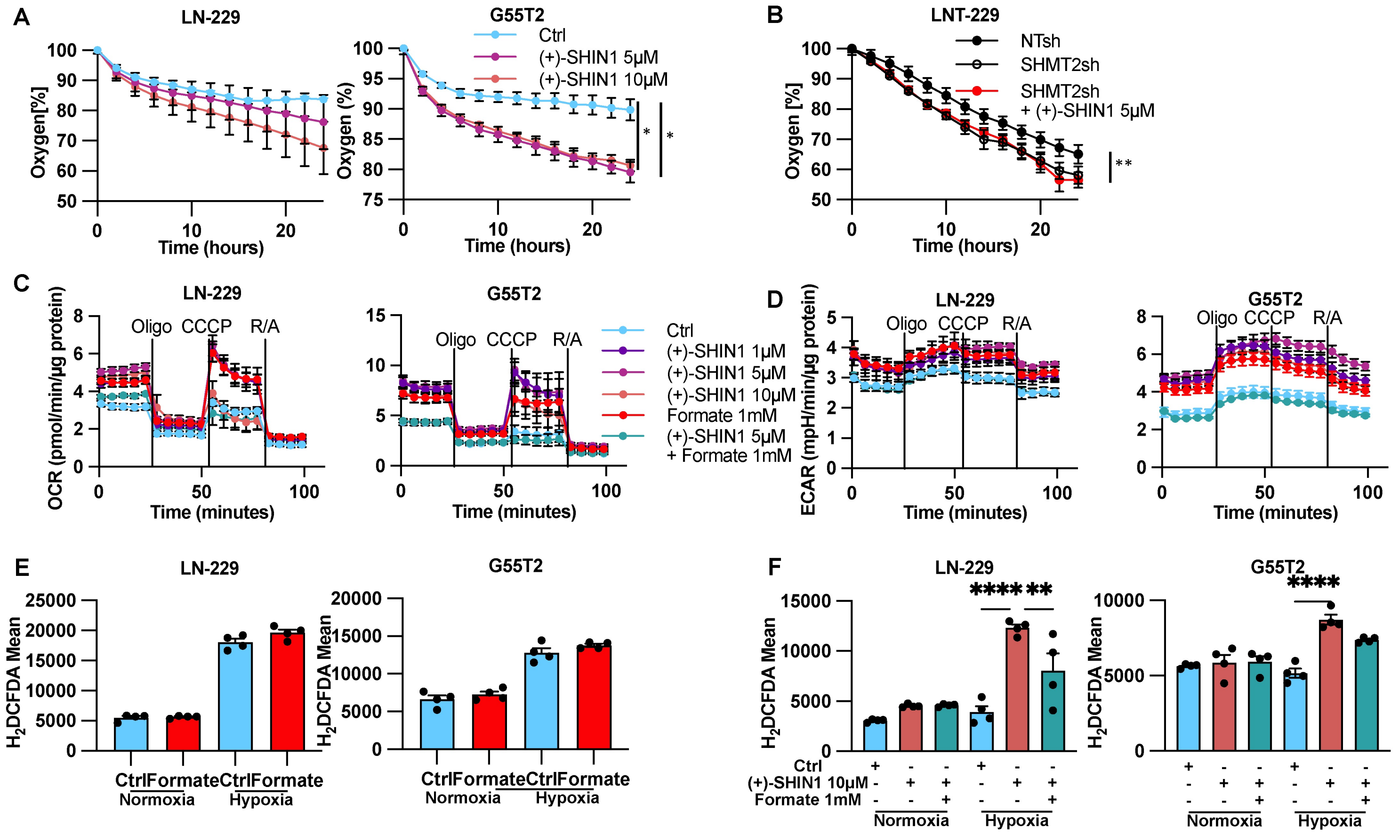
A
LN-229
Oxygen[%]
Time (hours)
G55T2
Oxygen (%)
Ctrl
(+)-SHIN1 5µM
(+)-SHIN1 10µM
B
LNT-229
NTsh
SHMT2sh
SHMT2sh
+ (+)-SHIN1 5µM
Oxygen [%]
C
OCR (pmol/min/µg protein)
Oligo
CCCP
R/A
Time (minutes)
(+)-SHIN1 1µM
Formate 1mM
(+)-SHIN1 5µM
+ Formate 1mM
D
ECAR (mpH/min/µg protein)
E
H2DCFDA Mean
Ctrl
Formate
Normoxia
Hypoxia
F
(+)-SHIN1 10µM
Formate 1mM

A

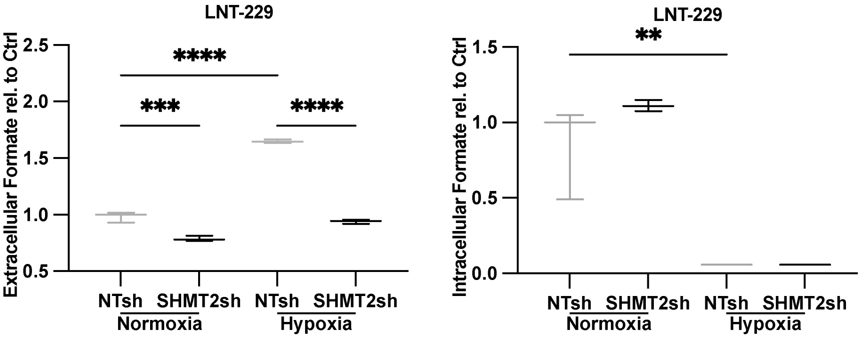

LNT-229
Extracellular Formate rel. to Ctrl
2.5
2.0
1.5
1.0
0.5
****
***
****
NTsh
SHMT2sh
NTsh
SHMT2sh
Normoxia
Hypoxia
LNT-229
Intracellular Formate rel. to Ctrl
1.5
1.0
0.5
0.0
**
NTsh
SHMT2sh
NTsh
SHMT2sh
Normoxia
Hypoxia

A

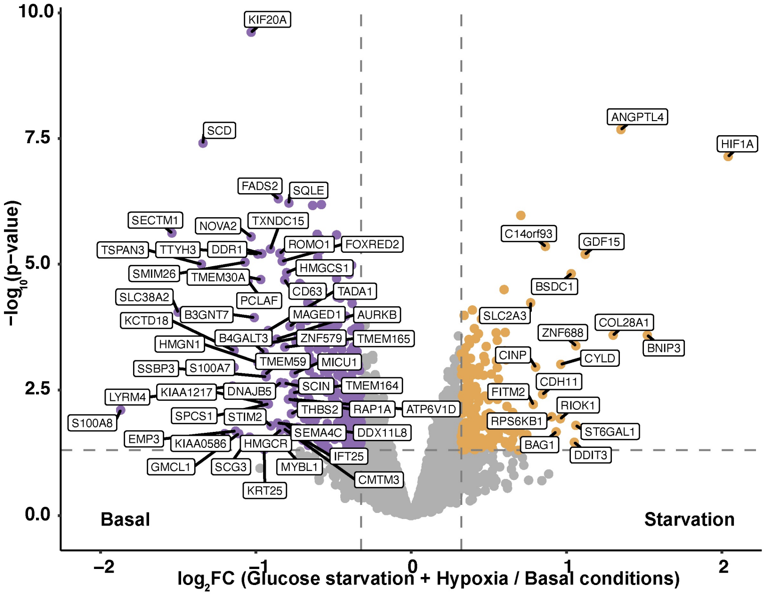

10.0
7.5
5.0
2.5
0.0
−log10(p−value)
−2
−1
0
1
2
log2FC (Glucose starvation + Hypoxia / Basal conditions)
Basal
Starvation
KIF20A
SCD
FADS2
SQLE
SECTM1
NOVA2
TXNDC15
TSPAN3
TTYH3
DDR1
ROMO1
FOXRED2
HMGCS1
SMIM26
TMEM30A
PCLAF
CD63
TADA1
SLC38A2
B3GNT7
MAGED1
AURKB
KCTD18
B4GALT3
ZNF579
TMEM165
HMGN1
TMEM59
MICU1
SSBP3
S100A7
SCIN
TMEM164
LYRM4
KIAA1217
DNAJB5
THBS2
RAP1A
ATP6V1D
S100A8
SPCS1
STIM2
SEMA4C
DDX11L8
EMP3
KIAA0586
HMGCR
IFT25
GMCL1
SCG3
MYBL1
CMTM3
KRT25
ANGPTL4
HIF1A
C14orf93
GDF15
BSDC1
SLC2A3
ZNF688
COL28A1
BNIP3
CINP
CYLD
CDH11
FITM2
RIOK1
RPS6KB1
ST6GAL1
BAG1
DDIT3

A (Fig. 2E, LNT-229)

**SHMT1**

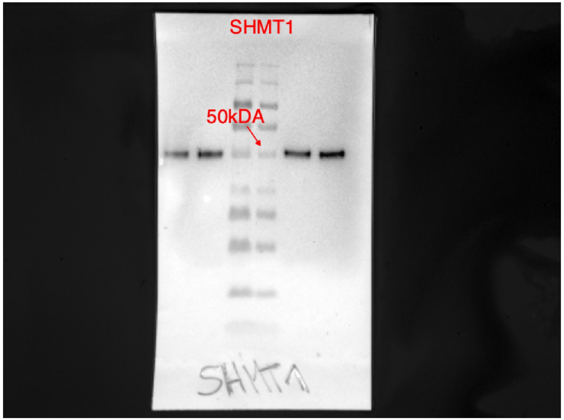


**Actin**

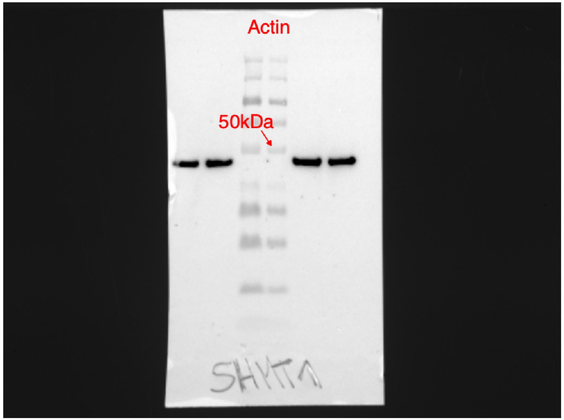


**SHMT2**

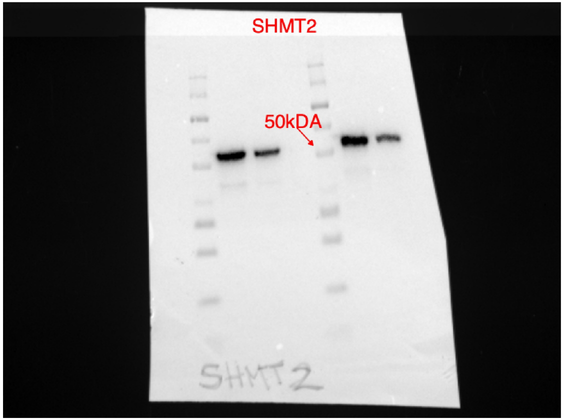


**Actin**

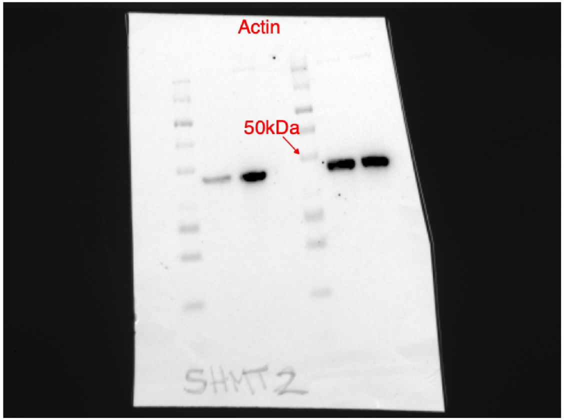

### Supplementary Figure legends

**Supplementary Figure 1**

**(A)** Area under the curve (AUC) values following (+)-SHIN1 and dimethyl sulfoxide (DMSO) treatment in serum-free DMEM containing 25 mM glucose in normoxia (21% $O_2$) in glioma cell lines LN-229, G55T2, NCH644 and patient-derived glioma stem cells BG7 (*n* = 3; mean with maximum and minimum). **(B)** Lentiviral SHMT2 knockdown was performed in glioma cell line LNT-229. SHMT1 and SHMT2 protein levels were analysed by immunoblot and normalised to β-actin, which served as loading control. **(C)** LN-229 and G55T2 cells were treated with 1 µM, 5 µM or 10 µM (+)-SHIN1 in serum-free DMEM containing 2 mM glucose in normoxia (21% $O_2$) or hypoxia (0.1% $O_2$) for 34 h. Cell death was analysed by lactate-dehydrogenase (LDH)-release assay (*n* = 4; mean ± standard error of the mean; one-way ANOVA followed by Tukey's multiple comparisons test; $^{**}P < 0.01$, $^{****}P < 0.0001$). **(D)** Cell viability of NCH644 and BG7 cells following (+)-SHIN1 treatment ± 1 mM formate in serum-free DMEM containing 25 mM glucose in normoxia (21% $O_2$) using the CellTiter-Glo assay (*n* = 3; mean ± standard error of the mean). **(E)** LN-229 and G55T2 cells were seeded (1,000 cells per well) in 6-well-plates, treated with 5 µM or 10 µM (+)-SHIN1 ± 1 mM formate in DMEM containing 10% FCS in normoxia (21% $O_2$) after 24 h and fixed by crystal-violet staining after 6 days (*n* = 3; mean; one-way ANOVA followed by Tukey's multiple comparisons test; $^{*}P < 0.05$, $^{**}P < 0.01$, $^{***}P < 0.001$, $^{****}P < 0.0001$). **(F)-(G)** LN-229, G55T2 and BG7 cells were treated with 10 µM (+)-SHIN1 ± 1 mM formate or 1mM formate in serum-free DMEM containing 2 mM glucose in normoxia (21% $O_2$) or hypoxia (0.1% $O_2$) for 22-34 h. Cell death was quantified by propidium-iodide staining (*n* = 4; mean ± standard error of the mean; one-way ANOVA followed by Tukey's multiple comparisons test; $^{**}P < 0.01$, $^{****}P < 0.0001$). Vehicle controls received DMSO at the concentration present in the highest (+)-SHIN1 dose.

**Supplementary Figure 2**

**(A)** Quantification of specific growth constant of the bulk tumour in the co-culture assay treated with 10 µM (+)-SHIN1 in mTeSR1-based medium with FGF2 and EGF (20 ng/ml each) in normoxia (21% $O_2$) for 3 days (*n* = 6; mean ± standard error of the mean; Welch's t-test; $^{*}P < 0.05$). **(B)** Quantification of specific growth constant of the embryoid body (EB) in the co-culture assay (*n* = 6; mean ± standard error of the mean; Welch's t-test). Representative images of the Fiji/ImageJ mask used for automated quantification of the tumour area in the threedimensional co-culture assay. Scale bar = 100 µm. Vehicle controls received DMSO at the concentration present in the highest (+)-SHIN1 dose.

**Supplementary Figure 3**

**(A)-(B)** Oxygen consumption of LN-229 and G55T2 treated with 5 µM or 10 µM (+)-SHIN1 and LNT-229 non-targeting shRNA (NTsh) and SHMT2-targeting shRNA (SHMT2sh) cells ± 5 µM (+)-SHIN1 was measured in DMEM containing 10% FCS in normoxia (21% $O_2$) overlaid with paraffin oil with a fluorescence-based assay. Oxygen consumption is shown relative to the start of the experiment ($n$ = 3; mean ± standard error of the mean; Welch's t-test of linear regression slopes; *$P$ < 0.05, **$P$ < 0.01). **(C)-(D)** LN-229 and G55T2 cells were treated with 1 µM, 5 µM or 10 µM (+)-SHIN1 ± 1 mM formate in serum-free DMEM in normoxia (21% $O_2$) for 24 h. After that oxygen consumption rate (OCR) and extracellular acidification rate (ECAR) were measured in the Seahorse Bioscience Extracellular Flux Analyzer XF96 and XF$^e$96 with port injections of oligomycin (Oligo), Carbonyl cyanide m-chlorophenyl hydrazone (CCCP), rotenone (R) and antimycin A (A) ($n$ = 3; mean ± standard error of the mean). **(E)-(F)** LN-229 and G55T2 cells were treated with 10 µM (+)-SHIN1 ± 1 mM formate or 1 mM formate in serum-free DMEM containing 2 mM glucose in normoxia (21% $O_2$) or hypoxia (0.1% $O_2$) for 18-24 h. Cells were analysed by 2′,7′-dichlorodihydrofluorescein diacetate (H2DCFDA) and PI staining. Only PI negative (= living) cells were quantified for H2DCFDA flow cytometry analysis ($n$ = 4; mean ± standard error of the mean; one-way ANOVA followed by Tukey's multiple comparisons test; *$P$ < 0.05, **$P$ < 0.01, ****$P$ < 0.0001). Vehicle controls received DMSO at the concentration present in the highest (+)-SHIN1 dose.

**Supplementary Figure 4**

**(A)** Extra- and intracellular formate levels in LNT-229 non-targeting shRNA (NTsh) and SHMT2-targeting shRNA (SHMT2sh) cells incubated in serum-free DMEM containing 25 mM glucose in normoxia (21% $O_2$) or 2 mM glucose in hypoxia (0.1% $O_2$) for 8 h ($n$ = 3; median with maximum and minimum; one-way ANOVA followed by Tukey's multiple-comparisons test; **$P$ < 0.01, ***$P$ < 0.001, ****$P$ < 0.0001).

**Supplementary Figure 5**

(**A**) Volcano plot of protein abundance in vehicle-treated LN-229 cells in serum-free DMEM containing 25 mM glucose in normoxia (21% $O_2$) versus 2 mM glucose in hypoxia (0.1% $O_2$) after 8 h, analysed by tandem mass tag-based mass spectrometry. Dashed lines indicate the thresholds applied for significance (|$\log_2$ fold change| = 0.32, corresponding to a fold change of 1.25, and $P = 0.05$); the y-axis shows $P$ values derived by limma[36,56] ($n = 3$).

### Supplementary methods

#### Analysis of public glioblastoma datasets

Dataset versions correspond to the latest releases available at the time of analysis. The CGGA data were retrieved from http://www.cgga.org.cn and the Ivy GAP data from https://glioblastoma.alleninstitute.org.

#### Cell lines and culture conditions

NCH644 and BG7 cells were cultured as suspension tumour spheres in NeuroBasal medium (Thermo Fisher Scientific, Germany) supplemented with 1% v/v GlutaMAX (Thermo Fisher Scientific, Germany), 2% v/v B-27 supplement (Thermo Fisher Scientific, Germany), 1% v/v penicillin/streptomycin (Thermo Fisher Scientific, Germany), 20 ng/ml recombinant human epidermal growth factor (EGF; PeproTech, USA) and 20 ng/ml recombinant human basic fibroblast growth factor 2 (FGF2; PeproTech, USA). When tumour spheres reached approximately 300 µm in diameter, they were dissociated into a single-cell suspension by incubation with Accutase (Sigma-Aldrich, USA) for 10 min at 37°C. Dissociated cells were pelleted, washed once with fresh medium and reseeded at $1 \times 10^5$ to $2 \times 10^5$ cells/ml in fresh culture medium.

#### Generation of fluorescent BG7 cells

For stable fluorescent labelling, BG7 cells were transduced with lentiviral vectors encoding turboGFP using the GIPZ lentiviral vector (Open Biosystems, USA). Lentiviral particles were produced as previously described (modified from Brown et al. 2020). Briefly, HEK293T cells were co-transfected with the respective transfer vector, the packaging plasmid pCMVdeltaR8.74 and the vesicular stomatitis virus glycoprotein envelope plasmid pMD.G.2 using calcium phosphate transfection with 2x HEPES-buffered saline (Thermo Fisher Scientific, Germany). Viral supernatants were harvested 72 h after transfection and used for transduction. Successfully transduced cells were selected with puromycin (1 µg/ml).

#### SHMT2 knockdown in LNT-229 cells

Cells were transfected according to the manufacturer's instructions using a lipophilic transfection reagent. After 24 h, medium was replaced and cells were cultured for an additional 48 h before puromycin selection. Single-cell clones were expanded and SHMT2 knockdown efficiency was validated by immunoblotting.

#### Cell density adjustment

To ensure comparable starting densities between sublines, cells were seeded at 100, 120 and 140% of the regularly used starting cell numbers. Following medium change, cells were fixed and stained with 0.5% crystal violet, extensively washed, and the dye was eluted in sodium citrate solution. Absorbance at 595 nm was measured in a microplate reader and used to normalise seeding densities across sublines.

#### Cell viability assay

Adherent cell lines were seeded at $5 \times 10^3$ cells per well in 96-well plates 24 h prior to treatment. Non-adherent spheroid cultures were seeded at $1.5 \times 10^3$ cells per well in ultra-low attachment U-bottom 96-well plates to allow spheroid formation. After 96 h of treatment, luminescence was measured according to the manufacturer's protocol using a multimode plate reader.

#### LDH release assay

Cells ($2 \times 10^4$ per well) were seeded in 96-well plates, treated under normoxic or hypoxic and nutrient-restricted conditions, and processed according to the manufacturer's instructions. Supernatants were collected after brief centrifugation and the remaining cells were lysed to determine maximal LDH. Absorbance at 490 nm was measured, the background ("low control") was subtracted, and relative LDH release was calculated as the ratio of supernatant to maximal LDH.

#### Flow cytometric assays

For propidium iodide (PI)-based viability assays, $1.2 \times 10^5$ cells per well were seeded in 24-well plates and treated as indicated. Supernatants were collected, adherent cells were washed with PBS, detached with trypsin, combined with the supernatants, washed and resuspended in flow cytometry buffer (PBS with 2% FCS). For H2DCFDA and MitoTracker staining, cells were harvested as above, incubated with the respective dye for 30 min at 37°C, washed and counterstained with PI. PI-positive cells were scored as non-viable, and the mean fluorescence intensity (MFI) of the PI-negative population was used as an indicator of intracellular oxidative stress.

#### Image analysis of invasion and migration assays

Transwell inserts were coated with diluted Matrigel and incubated for 2 h at 37°C for polymerisation. After removal of excess matrix, $2 \times 10^4$ cells in serum-free medium containing the respective treatments were added to the upper chamber. After 24 h, inserts were stained with crystal violet, non-invading cells on the upper surface were removed with cotton swabs, and membranes were excised, mounted and dried. Invaded cells were quantified using a standardised Fiji macro comprising automatic thresholding, binarisation, watershed separation and particle counting with fixed size and circularity criteria. For the gap-closure assay, LN-229 cells ($2 \times 10^4$ per chamber) were seeded in 70 µl medium and cultured overnight to form confluent monolayers. Inserts were removed, cells were washed with PBS, pre-treated with mitomycin C (10 µg/ml, 1 h), washed twice with PBS and incubated in serum-free medium. Gap areas were imaged at 0, 24 and 48 h using phasecontrast and live-cell imaging (4x objective) and quantified in Fiji by threshold-based segmentation.

#### Immunoblotting

Cells ($1.2 \times 10^6$) were seeded in 6-cm dishes and treated as indicated. Cells were washed twice with ice-cold PBS, snap-frozen and lysed in buffer containing 50 mM Tris-HCl pH 7.4, 150 mM NaCl, 1% NP-40, 0.5% sodium deoxycholate, 0.1% sodium dodecyl sulphate (SDS) and protease/phosphatase inhibitors. Protein concentrations were determined by Bradford assay. Membranes were blocked in 5% bovine serum albumin in Tris-buffered saline with Tween 20 and incubated overnight at 4°C with primary antibodies, followed by horseradish peroxidase-conjugated secondary anti-rabbit antibodies (Cell Signaling, Germany). For reprobing, membranes were stripped, reblocked and incubated with anti-beta-actin. Densitometry was corrected for background.

#### Oxygen consumption and extracellular flux measurements

For OxoDish measurements, cells ($1.2 \times 10^5$ per well) were seeded and cultured in 600 µl complete medium containing the indicated treatments. Wells were overlaid with sterile paraffin oil to minimise gas exchange with ambient air, and oxygen partial pressure was recorded every 15 min over 24 h using an optical sensor reader. For Seahorse measurements, cells ($3 \times 10^4$ per well) were seeded in XF96 plates, allowed to adhere and treated in serum-free DMEM for 24 h. Immediately prior to measurement, medium was replaced by bicarbonate-free, serum-free XF DMEM (Agilent, pH 7.4) supplemented with 25 mM glucose and 2 mM glutamine. Data were analysed with Wave software and normalised to the protein content of the wells.

#### Redox and enzyme activity assays

For the NADPH/NADP+ assay, cells ($2 \times 10^4$ per well) were seeded in 96-well plates, treated for 8 h and lysed in 0.2 N NaOH containing 1% dodecyltrimethylammonium bromide. NADPH and NADP+ were measured separately following sequential neutralisation and heat steps according to the kit protocol. For glutathione measurements, cells ($2 \times 10^4$ per well) were treated for 8 h and lysed according to the manufacturer's instructions; reduced (GSH) and oxidised (GSSG) glutathione and total glutathione were calculated from standard curves. For formate quantification, cells ($1 \times 10^6$ per well) were seeded in 6-well plates and treated for 8 h, medium supernatants were collected, and cells were washed with ice-cold PBS and homogenised in formate assay buffer by sonication ($1 \times 10^6$ cells per 100 µl buffer). Lysates were clarified by centrifugation (14,000 × g, 10 min, 4°C) and formate was quantified at 450 nm after enzymatic reaction at 37°C. For pyruvate kinase activity, cells ($1 \times 10^6$) were treated for 8 h, lysed on ice in four volumes of assay buffer and clarified by centrifugation (12,000 × g, 10 min, 4°C); absorbance at 570 nm was measured at baseline and after 20 min at room temperature in the dark, and activity was calculated from the change in absorbance using a pyruvate standard curve and normalised to cell number.

### Metabolomics

For the extraction of intracellular metabolites, cells were washed with ice-cold PBS, and 600 µL of pre-cooled 80% methanol (−80 °C) was added to each well. Cells were scraped on dry ice, incubated on dry ice for 15 min, and subjected to three freeze–thaw cycles at −80 °C. The resulting lysates were centrifuged at 15,000 rpm for 10 min at 4 °C and the supernatants were collected and stored at −80 °C. For semi-quantitative profiling of intracellular sugar phosphates, amino acids, TCA cycle intermediates, and energy metabolites, including AMP and ADP, supernatants were thawed on ice, vortexed, and centrifuged for 10 min at 4 °C. An aliquot of 60 µL from each supernatant was transferred to a 96-well plate for LC-QTOF-MS analysis. A pooled quality-control (QC) sample was prepared by combining aliquots from all samples and analyzed after every five samples to monitor analytical precision. The coefficient of variation was below 15% for all analytes. LC-QTOF-MS analysis was performed as described,[29] with the following modifications. Mobile phase A consisted of 10 mM ammonium acetate in water, adjusted to pH 7 and supplemented with 5 µM InfinityLab Deactivator Additive (Agilent Technologies, Germany). Mobile phase B consisted of 90% acetonitrile supplemented with 2.5 µM InfinityLab Deactivator Additive. Targeted feature extraction was performed using MassHunter Profinder (version B.08.00, Agilent Technologies). Metabolites were annotated by matching accurate masses and retention times to those of reference compounds analyzed alongside the samples. Peak areas were manually integrated and exported as comma-separated values (CSV) files for subsequent normalization and visualization. For extracellular lactate quantification, cell culture medium samples were diluted 20-fold with methanol (1:1, v/v) and analyzed by targeted LC-MS/MS metabolomics as described.[28]

### Metabolomic 13$C_6$-Flux analysis into polar metabolites

Incorporation of $^{13}C_6$-glucose into polar metabolites was examined using an UHPLC-HRMS screening method, as previously described[30] and as described in detail in the Supplementary Analytical protocol file (Excel). Briefly, 300,000 cells were suspended in 50 µl 1x phosphate buffered saline (1x PBS) and subjected to liquid-liquid extraction with 75 µl of internal-standard solution in methanol and 250 µl methyl tert-butyl ether. Following re-extraction of the upper phase, the lower phase was dried under nitrogen at 45 °C and stored at -80 °C until analysis. Samples were reconstituted in a 50:50 (v/v) mixture of acetonitrile and water. Two sets of quality control (QC) samples were prepared by pooling aliquots of the study samples and non-study plasma QC samples, respectively. Analysis was performed using a Vanquish Horizon UHPLC system coupled to an Orbitrap Exploris 480 mass spectrometer (both Thermo Fisher Scientific, Dreieich, Germany) operated in both positive- and negative-ionization modes. Chromatographic separation was achieved using a SeQuant ZIC-HILIC column (3.5 µm, 100 mm × 2.1 mm internal diameter; Merck, Darmstadt, Germany), equipped with a guard column of the same stationary phase and an inline filter. Gradient elution was performed using solvent A (0.1% formic acid in water) and solvent B (0.1% formic acid in acetonitrile). Data were acquired using Xcalibur version 4.4. Labeled and unlabeled metabolites were identified using Compound Discoverer version 3.3 and the mzCloud library (both Thermo Fisher Scientific, Dreieich, Germany). Identifications were manually reviewed, and duplicate features, e.g. those detected in both ionization modes, were removed, retaining features with higher signal intensities, fewer missing values, and better QC precision. Median-based probabilistic quotient normalization factors were calculated for each sample from Compound Discoverer peak areas across all detected ion features using a custom R pipeline. Up to three dominant exchanges, for which the labeled feature accounted for at least 10% of the signal in at least one sample group, were subsequently curated in Skyline version 26.1.0.266.[31] Normalized and raw feature peak areas (AUC = area under the curve) were exported as csv file for visualization and analysis of treatment effects. AUC were divided by 1000 to reduce large numbers and square root transformed to adjust a skewed distribution. For visualization as heatmaps, sqrt-AUC data were scaled to have a common median and variance of 1, referred to as Z-scores (autoscaling in MetaboAnalyst) according to the formula Z = (x – median)/SD, where x is the sample's value, and SD the standard deviation. Data were submitted to multivariate ANOVA for "treatment" by "time" and subsequent post-hoc analyses between treatments.

### Proteomics

#### MS sample preparation

For proteome analysis, 1 × $10^6$ cells per well were seeded in 6-well plates, treated as indicated, washed three times with ice-cold PBS, scraped in PBS and transferred to LoBind tubes. After centrifugation (15,000 rpm, 10 min, 4 °C), supernatants were removed and pellets were snap-frozen in liquid nitrogen and stored at −80 °C. Cells were lysed in Lysis Buffer (2% SDS, 50mM Tris pH 8.5, 10mM TCEP, 40mM CAA, supplemented with protease inhibitor cocktail and phosphatase inhibitors) sonicated at 4°C in a Bioruptor Pico2 (30/30, 10 cycles) and boiled at 95 °C for 10 min. Proteins were precipitated using methanol-chloroform and resuspended in 8 M urea, 50 mM Tris pH 8.5. Proteins were digested with 1:50 w/w LysC (Wako Chemicals) and 1:100 w/w trypsin (Promega) overnight at 37 °C after dilution to a final urea concentration of 1 M using 50 mM Tris pH 8.5. Digested peptides were then acidified (pH 2–3) using trifluoroacetic acid (TFA) and purified using C18 SepPak columns (Waters). Desalted peptides were dried and resuspended in TMT-labeling buffer (200 mM EPPS pH 8.2, 20% acetonitrile). 10µg of peptides per condition were subjected to TMT labeling with 1:2.5 peptide-to-TMT ratio (w/w) for 1 h at room temperature. The labeling reaction was quenched by addition of hydroxylamine to a final concentration of 0.5% and incubation at room temperature for 15 min. Successful TMT labeling was verified by mixing equimolar ratios of peptides and subjecting the mix to single shot LC-MS/MS analysis. Peptides were fractionated using the high pH reverse-phase fractionation kit (Thermo Fisher Scientific) into 16 fractions and afterwards concatenated into 8 fractions. The pooled fractions were dried in a vacuum concentrator and resuspended in 2% ACN, 0.1% TFA for LC-MS analysis.

#### Mass spectrometry data acquisition for proteomics

Tryptic peptides were analyzed on an Orbitrap Fusion Lumos coupled to an easy nLC 1200 (Thermo Fisher Scientific) using a 25 cm long, 75µm ID fused-silica column packed in house with 1.9 µm C18 particles (Reprosil pur, Dr. Maisch), and kept at 50°C using an integrated column oven (Sonation). HPLC solvents consisted of 0.1% Formic acid in water (Buffer A) and 0.1% Formic acid, 80% acetonitrile in water (Buffer B). The peptides were eluted by a non-linear gradient from 3% to 24% Buffer B over 150 minutes, followed by a stepwise increase to 90% Buffer B in 6 minutes, which was held for another 9 minutes. A synchronous precursor selection (SPS) multi-notch MS3 method was used in order to minimize ratio compression as previously described.[32] Full scan MS spectra (350-1400 m/z) were acquired with a resolution of 120,000 at m/z 200, a maximum injection time of 100 ms, and an AGC target value of 4 x 105. The 10 most intense precursors with a charge state between 2 and 5 per full scan were selected for fragmentation ("Top Speed" with a cycle time of 1.5 seconds) and isolated with a quadrupole isolation window of 0.7 Th. MS2 scans were performed in the Ion trap (Turbo) using a maximum injection time of 50 ms, an AGC target value of 15000, and fragmented using CID with a normalized collision energy (NCE) of 35%. MS3 acquisition was performed on the 10 most intense MS2 fragment ions with an isolation window of 0.7 Th (MS) and 2 m/z (MS2). Ions were fragmented using HCD with an NCE of 50% and analyzed in the Orbitrap with a resolution of 50,000 at m/z 200, a scan range of 110-500 m/z, an AGC target value of 150000 and a maximum injection time of 86 ms. Repeated sequencing of already acquired precursors was limited by setting a dynamic exclusion of 60 seconds and 7 ppm, and advanced peak determination was deactivated. All spectra were acquired in centroid mode.

#### Mass spectrometry proteomic data analysis

MS raw data were analyzed using FragPipe v22.0, with MSFragger v.4.3[33] and Philosopher v.5.1.0.[34] Acquired spectra were searched against the human reference proteome (Taxonomy ID 9606) downloaded from UniProt (07/08/2024; 20,418 entries) with a precursor mass tolerance of 20 ppm and a fragment mass tolerance of 20 ppm. Identifications were filtered to obtain false discovery rates (FDR) below 1% for both peptide spectrum matches (minimum peptide length of 7) and proteins using a target-decoy strategy. For all searches, carbamidomethylated cysteine was set as a fixed modification and oxidation of methionine and N-terminal protein acetylation as variable modifications with allowing up to 3 modifications per peptide. Strict trypsin cleavage was set as protein digestion rule. Label-free quantification was performed using IonQuant v.1.11.11.[35] Data were further processed using FragPipe Analyst.[36]

#### Generation of embryoid bodies

Human induced pluripotent stem cells (DF-19-9-11T.H, WiCell Research Institute, Madison, WI, USA) were used for embryoid body formation following the STEMdiff Cerebral Organoid workflow (STEMCELL Technologies, Canada) with minor modifications. Cultures were used at approximately 70-80% confluency with minimal spontaneous differentiation. Cells were washed with D-PBS, dissociated with Accutase (Sigma-Aldrich), collected and centrifuged at 300 × g for 5 min. Cells were resuspended in Organoid Formation Medium supplemented with 10 µM Y-27632 (STEMCELL Technologies) and adjusted to $9 \times 10^4$ cells/ml. For embryoid body formation, 100 µl of the suspension, corresponding to $9 \times 10^3$ cells, was seeded per well into 96-well U-bottom ultra-low attachment plates (SBio, USA). Plates were incubated at 37°C and 5% CO2 and left undisturbed for the first 24 h to allow aggregate formation. On days 2 and 4, 100 µl of Organoid Formation Medium was added gently to each well. Embryoid bodies were cultured until day 5, by which time they reached approximately 500 µm in diameter and displayed a compact, round morphology with smooth edges.[37]
